# Dynamic profiling of the aggresome processing pathway

**DOI:** 10.1101/2025.10.30.684516

**Authors:** Sucheta Ghosh, Longlong Wang, Daniel Hess, Vytautas Iesmantavicius, Jan Englinger, Chun Cao, Gabriele Matthias, Jan Seebacher, Patrick Matthias

**Affiliations:** Friedrich Miescher Institute for Biomedical Research (FMI); Basel 4056, Switzerland; Institute of Pharmaceutical Sciences, Department of Chemistry and Applied Biosciences, ETH Zurich; Zurich 8093, Switzerland; Faculty of Sciences, University of Basel; Basel 4031, Switzerland

**Keywords:** Aggresome processing pathway, stress response, HDAC6, Vimentin, proximity labeling, mass spectrometry

## Abstract

Cellular homeostasis is maintained by the efficient degradation of proteins by the 26S proteasome, a key component of the ubiquitin proteasome system. When the proteasome function is impaired due to cellular stress like mutations, chemicals or epigenetic stressors, there is accumulation of unfolded or misfolded, ubiquitinated proteins leading to cellular toxicity. Cells combat this rise in cellular toxicity by sequestering ubiquinated proteins in a transient cellular structure called the aggresome. The transport of this ubquitinated protein cargo along the microtubules to form aggresomes and their clearance from cells by autophagy is called the aggresome processing pathway (APP). APP activation is a hallmark of several neurodegenerative diseases and has also been implicated in Influenza A Virus uncoating step. To study the dynamics of APP and the compositional diversity of the aggresome, we used time-dependent proximity labeling, mass spectrometry, and immunofluorescence microscopy. Here, we used vimentin (VIM) and histone deacetylase 6 (HDAC6), two established aggresome markers as baits to visualize aggresome formation and clearance in HeLa cells. We identified more than 75 novel components of mature aggresome. Further, we characterised the time-dependent protein-protein interactions in the neighbourhood of VIM and HDAC6 during APP progression. Our study identified new interactors of VIM and HDAC6 that maybe involved in aggresome clearance by autophagy. Together, our findings uncover novel APP-associated proteins that are central to advancing the understanding of this poorly characterized and highly complex pathway implicated in multiple diseases.

## Introduction

Protein quality control is essential for healthy functioning of an organism. Several regulatory pathways work in collaboration to maintain protein homeostasis within a cell. Unfolded or misfolded proteins are degraded by the ubiqitin proteasome system (UPS) (Ciechanover & Schwartz, 1998). When the proteasome is overwhelmed due to genetic mutations, inappropriate protein assembly, aberrant modifications or environmental stress, these misfolded proteins have the propensity to form aggregates. In cells, when the UPS function is impaired, misfolded proteins tend to accumulate over time as they migrate towards the perinuclear region leading to formation of aggresomes near the microtubules-organizing center (MTOC) (Johnston et al., 1998). The aggresomes are then cleared via autophagy (Fortun et al., 2003; Klionsky et al., 2010). Therefore, the APP is a stress response pathway arising in cells stressed with overload of ubiquitinated and misfolded proteins due to an impaired proteasome. Little is known about the proteins involved in APP and about aggresome composition during proteasomal inhibition.

The cytosolic lysine deacetylase enzyme, HDAC6, and the well-known protein modifier ubiquitin (Ub) are some of the key players that have been identified in the APP. HDAC6 has a zinc-finger Ub binding domain (ZnF-UBP) on its C-terminus that is very important for the formation of cellular granules in response to cellular stress. It has been shown that in presence of a stress signal, HDAC6 recruits unanchored polyubiquitin (polyUb) chains via ZnF-UBP (Hook et al., 2002; Seigneurin-Berny et al., 2001). HDAC6 also indirectly interacts with the actin-myosin network (Artcibasova et al., 2023; Banerjee et al., 2014) and dynein, a motor protein that directs this complex of HDAC6-polyUb chains towards the aggresomal formation site (Kawaguchi et al., 2003). From a structural point of view, HDAC6 cannot recognize polyUb chains anchored to misfolded proteins because the Ub C terminal diglycine motif is not exposed. A reasonable molecular model to explain the physical interaction between HDAC6 and aggregated proteins has been proposed (Ouyang et al., 2012), which postulates that the deubiquitinase ataxin-3 is first recruited to aggregated proteins and partially cleaves the polyUb chains through its deubiquitination activity, this then allows HDAC6 to recognize aggregated proteins through the exposed and unanchored Ub C-termini. HDAC6 not only contributes to aggresome formation but also plays important roles for its clearance process by autophagy. It was reported that HDAC6 deacetylates cortactin which is recruited to protein aggregates where it enhances F-actin polymerization to clear aggregates by autophagy. POH1 (Proteasome non-ATPase subunit, regulatory subunit 11), a proteasomal deubiquitinase mediates release of unanchored Ub chains in aggresome that activate HDAC6 and promote aggresome clearance (Hao et al., 2013).

VIM, another aggresome marker plays a crucial role in aggresome biogenesis, acting as a cytoskeletal scaffold that facilitates the sequestration and transport of misfolded protein aggregates to the MTOC (Ivaska et al., 2007). Microscopy studies have shown that VIM forms a dense, cage-like structure around aggresomes, preventing the toxic dispersion of misfolded proteins into the cytoplasm and enhancing their subsequent clearance via autophagy (Garcia-Mata et al., 1999). HDAC6 binds to ubiquitinated misfolded proteins via its C-terminal ZnF-UBP and recruits them to the aggresome, while simultaneously interacting with VIM to mediate structural changes in the filament network (Kawaguchi et al., 2003).

This phenomenon of accumulation of misfolded proteins in proteinaceous inclusions like aggresomes has mostly been studied in the context of prominent pathological features common to many age-related neurodegenerative diseases like Parkinson’s disease, Alzheimer’s disease or Huntington’s disease (Chen et al., 2011). Previous studies from various labs including ours have shown that multiple stress pathways like formation of stress granules (Kwon et al., 2007) and aggresomes (Boyault et al., 2007), infections by influenza A virus (IAV) (Banerjee et al., 2014), or Zika virus (Wang et al., 2022) and development of inflammasomes (Magupalli et al., 2020) use similar components related to the APP. In particular, the aggresome/HDAC6 pathway can be hijacked by exogenous pathogens, like IAV to facilitate the infection. While some of the components have been identified, the precise understanding of their dynamics and interactions with other proteins in the network, is lacking. In this study, we have used a combination of acorbate peroxidase (APEX2) proximity labeling, mass spectrometry and immunofluorescence microscopy approach to significantly expand the repertoire of proteins involved in the APP during its progression and clearance. To study the dynamics of APP, we have used two aggresome markers, HDAC6 and VIM, and have established cell lines expressing them as fusion proteins to APEX2-GFP, allowing for mass spectrometry and immunofluorescence microscopy analysis. Through this approach we identified more than 75 previously unknown components of the mature aggresome. We report the existence of a highly integrated, dense interaction network of proteins in the neighbourhood of VIM and HDAC6 in unstressed cells. We have expanded the repertoire of proteins by identifying new interactors of VIM and HDAC6 that may play a role in aggresome clearance by autophagy. Therefore, our study can be used as a resource to understand the spatiotemporal landscape of mature aggresomes. It can be used as a starting point for further biochemical experiments to understand the molecular events during aggresome formation and clearance from cells. Collectively, our findings reveal novel components of the APP, offering pivotal insights into this poorly understood yet critically important pathway involved in the pathogenesis of multiple diseases.

## Results

### Dual-bait APEX2 proximity labeling allows for specific biotin labeling of aggresomes

To achieve a better understanding of the proteomic changes and compositional diversity of the APP during cellular stress, we have used proximity labeling proteomics and immunofluorescence microscopy to dissect and characterise this pathway (Fig.1A). We have used two well-known aggresome markers VIM (Johnston et al., 1998) and HDAC6 (Kawaguchi et al., 2003) as baits to obtain insights of the APP from a dual-bait viewpoint. We captured the interacting proteins in the neighbourhood of VIM and HDAC6 and monitored the changes in protein-protein interactions in unstressed cells and during aggresome biogenesis and clearance (Fig.1B). Firstly, we validated the APEX2 enzyme functionality by fusing APEX2 to HDAC6 and tested all conditions for optimum enrichment of biotinylated proteins from HDAC6 KO HEK293T cells (Saito et al., 2019) transiently transfected with an HDAC6-APEX2-GFP construct (Fig.S1A-C). Next, we generated VIM knockout (KO) and HDAC6 KO HeLa cells (Fig.S1D-E) and transduced the respective KO cells with VIM-APEX2-GFP and HDAC6-APEX2-GFP lentivirus constructs to establish stable cell lines. We validated the APEX2 activity and fusion protein expression of the established cell lines (Fig.1C-D and S1F-H). To induce proteasomal impairment, we used MG132, a classical proteasome inhibitor. We first validated aggresome formation in wild-type (WT) HeLa cells and treated them with MG132 or bortezomib, another proteasome inhibitor, for 18h hours; the treatment with both drugs induced clear aggresome formation, as detected by Ub staining and both aggresome markers VIM and HDAC6 localised to aggresomes (Fig.1E and S1I). We also validated by immunofluorescence microscopy that HDAC6 and VIM are essential for aggresome formation. After 18 hours MG132 exposure to VIM KO HeLa cells, no HDAC6-positive and Ub-positive aggresomes were detected as confirmed by HDAC6 and Ub antibody staining compared to WT HeLa cells (Fig.1F and S2A-B). Similarly, VIM-positive and Ub-positive aggresomes failed to form in HDAC6 KO HeLa cells post stimulation for 18 hours by MG132 as confirmed by VIM and Ub antibody staining compared to WT HeLa cells (Fig.1F and S2C-D). In conclusion, HDAC6 and VIM are essential for the cell to direct misfolded, aggregated proteins into aggresomes at the MTOC and for clearance of aggresomes by autophagy, in agreement with earlier studies (Hao et al., 2013; Johnston et al., 1998; Kawaguchi et al., 2003; Morrow et al., 2020). Next, we treated the established VIM-APEX2-GFP and HDAC6-APEX2-GFP cell lines with MG132 for 18 hours and visualized aggresome formation by monitoring GFP expression. These HDAC6- and VIM-APEX2-GFP cell lines allow robust and rapid biotin labeling of aggresome proteins in the presence of biotin-phenol (BP) and hydrogen peroxide (H_2_O_2_) (Fig.1G). As a specificity control, cells with constitutive expression of APEX2 localised in the cytoplasm (NES-APEX2-GFP) show a diffuse GFP signal and a biotinylation pattern that was unaffected by MG132.

**Figure 1:**
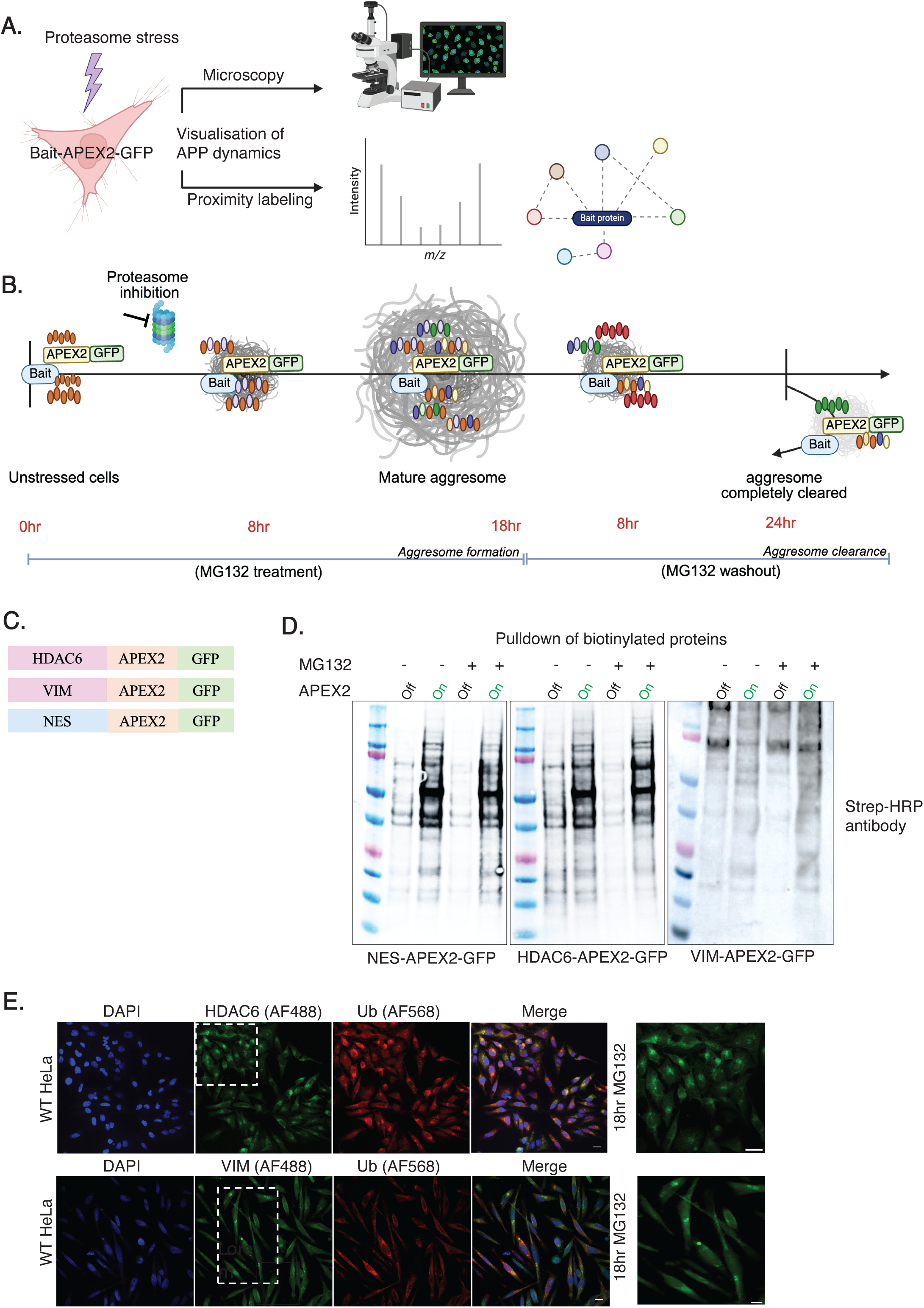
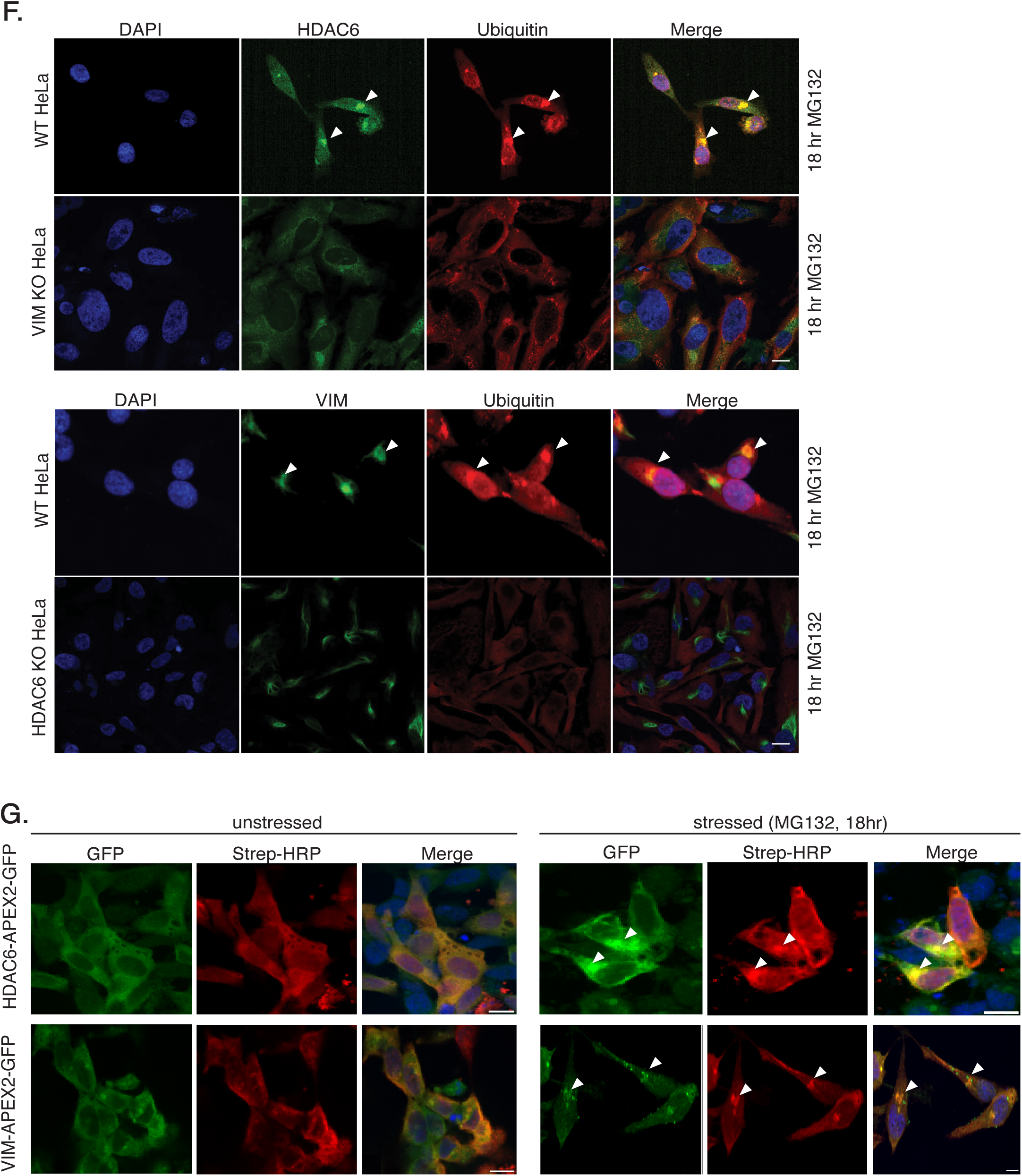
APEX2-GFP fused to HDAC6 or VIM localizes to the aggresome and biotinylates proteins efficiently upon proteasomal inhibition A) Schematic outlining the experimental design to study the aggresome processing pathway dynamics by APEX2-based proximity labelling and fluorescence microscopy. The established cell lines express bait (aggresome marker) proteins tagged with GFP and APEX2. The GFP tag enables to visualize the kinetics of APP. The APEX2 tag is used to study the interactome of the marker proteins at different time points of cellular stress. B) Schematic of HDAC6 and/or VIM interactome changes upon stress. This schematic explains our aim to capture the changes in interactome of VIM and HDAC6 during aggresome biogenesis and clearance in mammalian cells. C) Construct design of the C-terminal APEX2-GFP fusions to aggresome markers HDAC6 and VIM and of a control containing a nuclear export sequence (NES). These fusions were cloned into the lentiviral vector backbone and stably transduced into HDAC6 KO or VIM KO HeLa cells to generate stable APEX2 cell lines. D) Streptavidin-HRP immunoblot analysis of induced protein biotinylation in cell lysates from NES-APEX2-GFP, HDAC6-APEX2-GFP and VIM-APEX2-GFP HeLa cells that were untreated (control) or treated with MG132 for 18hr. On and Off refers to the biotinylation activity of APEX2 (APEX On-cells had been treated before lysis for 30 min with biotin-phenol and 2 min with H_2_O_2_, APEX Off-cells had been treated before lysis only 2 min with H_2_O_2_, BP treatment was omitted). E) Endogenous HDAC6, VIM and Ub localise to aggresomes when the proteasome is inhibited. HDAC6, VIM and Ub antibody staining of aggresomes in WT HeLa cells treated with MG132 treatment for 18hr to induce proteasome stress. Scale bar is 20μm. F) Immunofluorescence microscopy images indicating that VIM KO or HDAC6 KO leads to impairment of aggresome formation. WT HeLa, VIM KO and HDAC6 KO HeLa cells were treated with 10μM MG132 for 18hr. Aggresome formation (white arrowheads) was observed in WT HeLa cells validated by the overlap of HDAC6 or VIM signal with Ub signal *(panel 1 & 3)*. No HDAC6+ and Ub+ aggresomes were detected in VIM KO HeLa cells upon 18 hours proteasome inhibition *(panel 2).* Similarly, VIM+ and Ub+ aggresomes were absent in HDAC6 KO HeLa cells upon 18 hours proteasome inhibition *(panel 4)*. Scale bar is 20μm. G) HDAC6-APEX2-GFP and VIM-APEX2-GFP fusion proteins are active at the aggresome. Immunofluorescence microscopy images of streptavidin staining of unstressed cell show biotin signal dispersed in the cytoplasm. 18hr MG132-treated HDAC6-APEX2-GFP and VIM NES-APEX2-GFP HeLa cells show biotin signal localised to aggresomes (white arrowheads). The biotin signal colocalizes with the fusion-protein distribution in each condition as visible in the Merge image panel. Scale bar is 20μm.

### Microscopy based study of APP kinetics

We performed immunofluorescence microscopy at different time points of MG132 treatment and washout in cells to identify reference time points for our proximity labelling experiments. First, to identify the peak time of aggresome formation in HeLa cells, we treated WT cells with MG132 and analysed them at different times up to 18 hours (Fig.2A). We quantified Ub positive aggregates from 500 cells at each time point-the total number cells with Ub positive aggregates or aggresomes were plotted for each time point normalised to total number of cells quantified. We observed the occurrence of small Ub positive aggregates at 8 hours. This time point is referred to as “early aggresome formation”. These aggregates grew in size and we observed the maximum number of mature aggresomes in cells at 18 hours of proteasomal inhibition by MG132 (Fig.2B). Therefore, this time point is hereafter referred to as the “peak aggresome formation”. We next performed a similar analysis in the HDAC6-APEX2-GFP, VIM-APEX2-GFP and NES-APEX2-GFP cell lines and additionally included time points before and after peak aggresome formation (Fig.2C). The GFP signal from the fusion proteins in each of the two aggresome marker cell lines showed clear aggresome formation at 18 hours of MG132 treatment (Fig.2C, white arrows), that colocalised with the Ub staining, confirming that these structures are indeed aggresomes (Fig.S3A). As expected, the GFP signal from the NES-APEX2-GFP cell line at 18 hours of MG132 treatment was diffuse and did not allow for aggresome visualisation, but the Ub staining confirmed the presence of aggresomes in these cells (Fig.S3A). Similarly, the fusion protein from each cell line showed no aggregated GFP signal after 24 hours of MG132 washout post peak aggresome formation. Therefore, we refer to this time point as the “complete aggresome clearance”, with aggresomes being totally cleared from cells. The clearance of aggresomes was also confirmed by the strong decrease in the number of GFP positive aggresomes at 24 hours MG132 washout (Fig.2C). In addition, an intermediate time point at 8 hours of MG132 washout post 18 hours of MG132 treatment was examined by immunofluorescence microscopy and represents an “early time point of aggresome clearance”. At this stage, some cells still contain Ub-positive small aggregates as observed by immunofluorescence (Fig.S3A).

**Figure 2:**
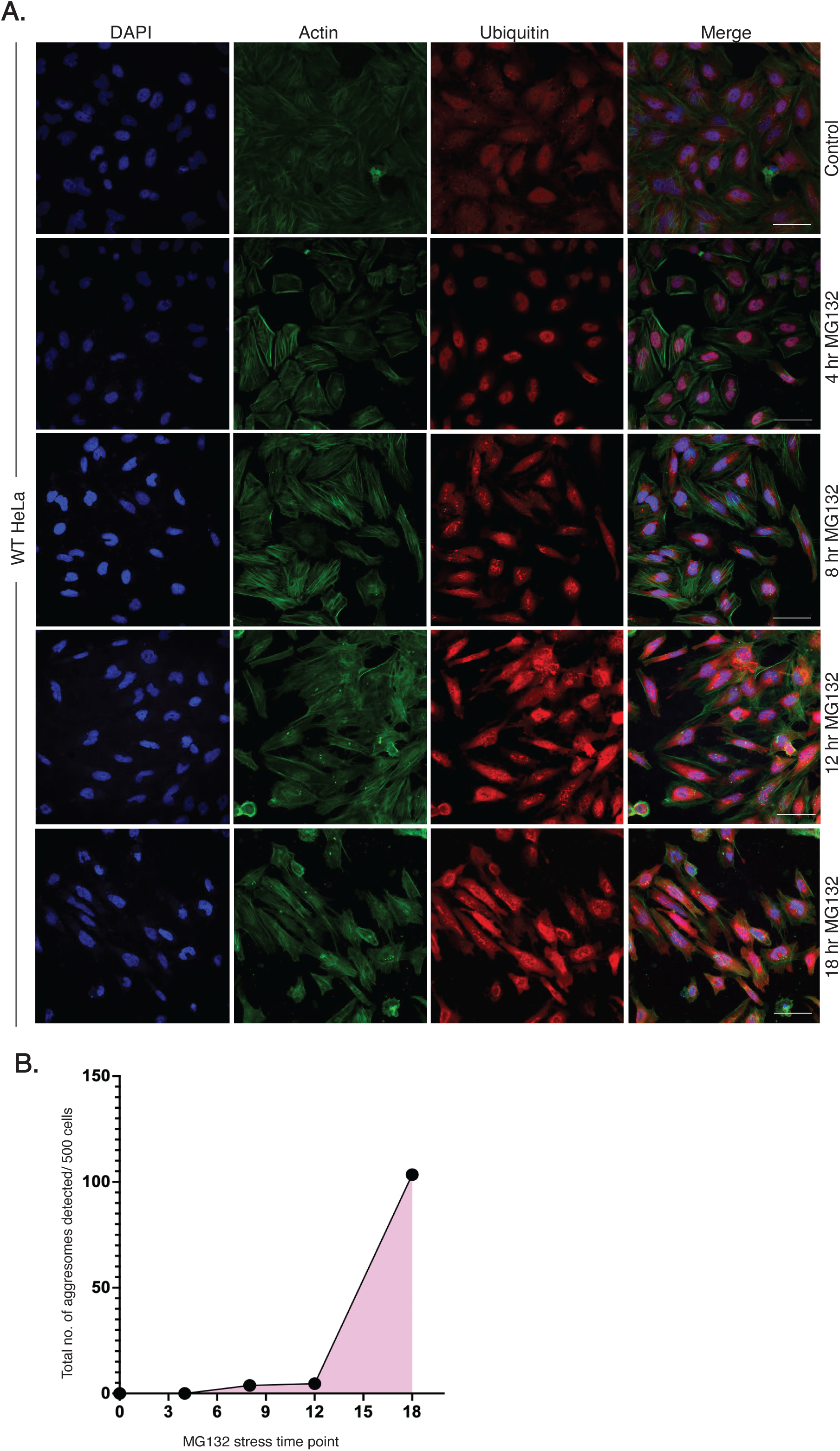
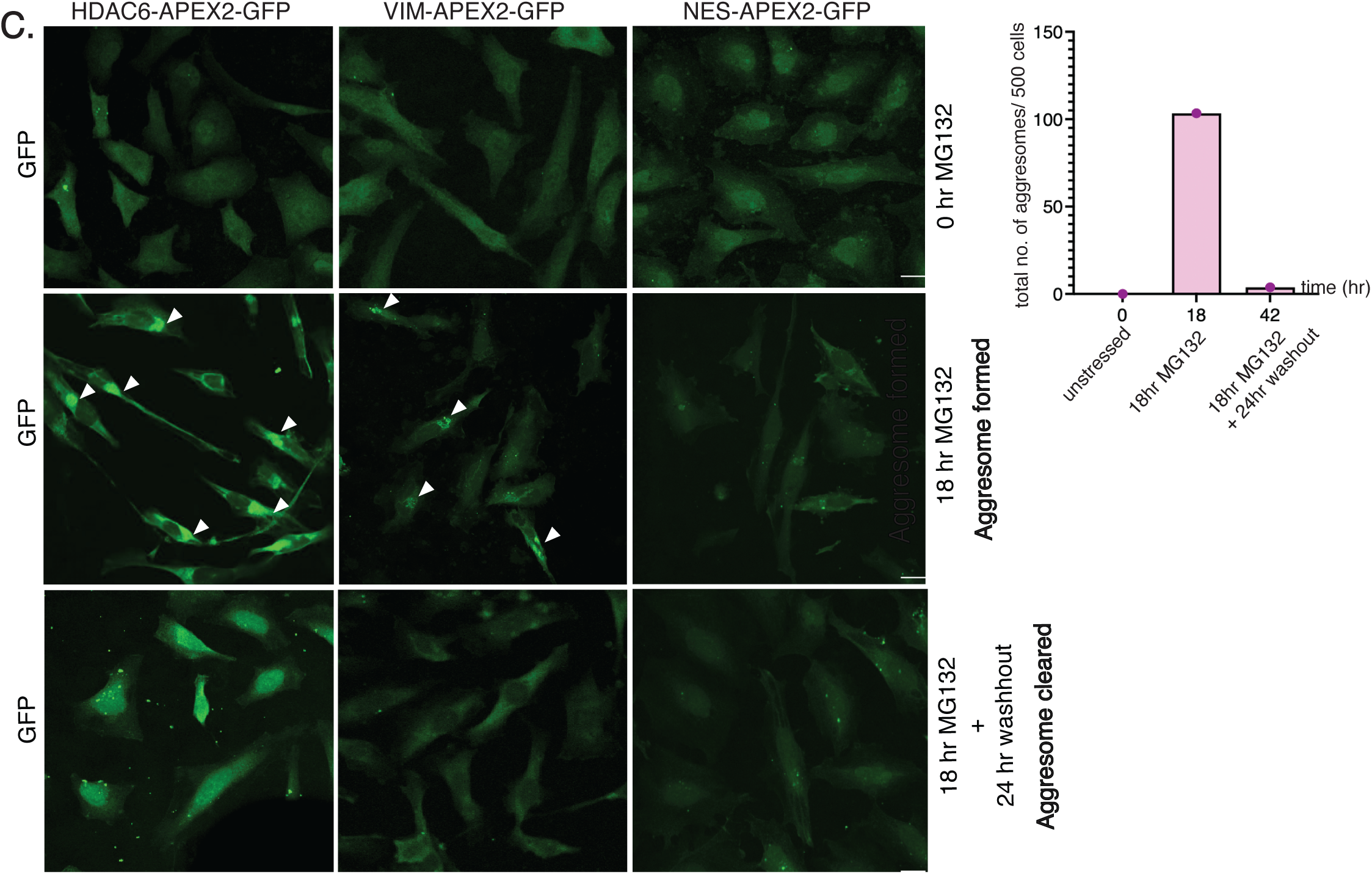
Kinetics of aggresome assembly in HeLa cells A) Immunofluorescence microscopy images of actin and Ub antibody staining of WT HeLa cells under MG132 treatment at different time points. Cells were untreated (control) or treated with 10μM MG132 for 4hr, 8hr, 12hr and 18hr. Clear aggresome formation was observed in cells under 18hr MG132 exposure. Scale bar is 50μm. B) Quantification of Ub^+^ aggregates in WT HeLa cells at different time points of MG132 treatment. For each condition, a total of 500 cells were analyzed per time point across multiple randomly selected fields to ensure representative sampling. Total number of Ub+ aggregate positive cells are plotted for each time point of MG132 treatment. C) Live GFP fluorescence signal (signal from 488nm laser of scanning disk confocal microscope) from fixed HDAC6-APEX2-GFP, VIM-APEX2-GFP HeLa cells, NES-APEX2-GFP cells showing mature aggresome formation at 18h MG132 treatment (white arrows point to aggresome) and clearance after 24h MG132 washout. Untreated cells were used as control. Scale bar is 20μm.

### Dual-bait proximity labeling reveals new aggresome proteins when the proteasome is inhibited

We aimed to understand the composition of mature aggresome at the MTOC in cells in which the proteasome is inhibited by MG132. We conducted APEX2 labeling experiments at selected time points of APP that were characterised earlier by immunofluorescence microscopy. At each time point of MG132 stress, VIM-APEX2-GFP, HDAC6-APEX2-GFP and NES-APEX2-GFP HeLa cell lines were treated with both substrates of APEX2, biotin phenol (BP) and hydrogen peroxide (H_2_O_2_, APEX On condition) along with samples to control for nonspecific streptavidin-bead binding (no BP, APEX Off condition) (Fig.S4A). Each labeling reaction was done in triplicate and streptavidin purification of biotinylated proteins was followed by label-free quantitative mass spectrometry. Here, we report the composition of the aggresome using the VIM-APEX2 bait at 18 hours MG132 exposure. We used a stringent significance cut off (log_2_ fold-change >-1 and <1 and adjusted p value=0.05) to identify proteins significantly above threshold (Fig.S4B). From this dataset, we employed a series of four comparison groups to systematically identify three classes of VIM-interacting proteins in stressed and unstressed cells: stress-dependent, stress-independent and stress-sensitive VIM interactors (Fig.3A). First, to identify stress-dependent VIM interactors, which associate with VIM only under proteasomal stress, we characterized biotinylated proteins in stressed versus unstressed VIM-APEX2-GFP cells (group 1). Next, we compared lysates from stressed VIM-APEX2-GFP cells incubated with BP and H_2_O_2_ (APEX2 On) to lysates of cells for which the BP substrate was omitted (APEX2 Off) (group 2). Third, to control for diffuse cytoplasmic labeling by VIM-APEX2-GFP, we also compared lysates from stressed VIM-APEX2-GFP and NES-APEX2-GFP cells (group 3). Last, to define stress-independent interactors as well as stress-sensitive VIM interactors, we profiled lysates from unstressed VIM-APEX2-GFP and NES-APEX2-GFP cells (group 4). We identified 3502 proteins across all samples using a conservative analysis pipeline (Fig.S4B), with very high correlation across all three technical replicates shown in the correlation plot for all samples (Fig.S5). In total, we detected 2814 proteins in at least 2 samples and 2606 proteins observed in at least 2 samples from the same condition. Protein identifications were highly reproducible across technical replicate experiments (Fig.S4C). As a control, we observed constitutively biotinylated proteins only enriched in APEX2 off samples (Fig.S4D). Several previously known well-characterized aggresome proteins were clearly enriched in our comparison of VIM-APEX2 hits versus NES-APEX2 hits at 18 hours of MG132 treatment. (e.g., BAG3, TUBB, HSPA1A, HSPA1B highlighted in bold black; Fig.3B, group 1 and 3). Known SG proteins were significantly enriched across all four comparison groups, with the greatest fold change detected in groups 1 and 3. To identify a group of aggresome proteins, a summation of significantly enriched proteins across all four comparison groups was performed. Although a total of 477 proteins were identified only in group 1, we identified 55 proteins shared between groups 1 and 3, 1 protein common to groups 2 and 3 and 19 proteins shared between groups 1, 2 and 3, as represented in the upset plot (Fig.3C). In total, we identified 75 new proteins that are interacting with VIM in the aggresome (Fig.S4E). Of these, four major classes of proteins were identified as new components of the mature aggresome: proteins involved in microtubular transport, stress granule components as well as regulators of autophagosome-lysosome fusion and autophagy, besides capturing known aggresome proteins like TUBB, HSPA1A, HSPA1B and VIM (Fig.3D). Our findings identify proteins involved in cellular localization and establishment of localisation as aggresome components (Fig.S4F). Among these 75 proteins, we hypothesize that enriched candidates like LAMP2, ABCD1, RAB9A, VAMP7, SCARB2, SNX12, SEC13, RER1, CALM3, MAPRE2, DDX19A, TWF2, ISG15, RANBP1, ARMCX3, CLTA, BAG3, NDUFAF2, SFN, HNRNPAB, MAP4, CPT1A and PPP6C support the VIM mediated transport of aggregated proteins towards the MTOC. Some other enriched proteins like ABCD1, ABCB7, ATP6V1G1 and LASP1 have transporter activity in cells. We identified a repertoire of APP-relevant proteins as VIM interactors in the aggresome. For example, the carrier protein, A2M has been shown to interact with aggregation-prone proteins, facilitating their internalization and degradation via the lysosomal pathway (Tomihari et al., 2023). Further, we identified ANXA11 as an aggresome component that is known to regulate protein homeostasis, particularly in the context of neurodegenerative diseases. Another major stress-dependent VIM aggresome interactor is ISG1 (Interferon-Stimulated Gene 15) which is a Ub-like protein that becomes conjugated to target proteins in response to interferon signaling, a process known as ISGylation. It was reported that the degradation of ISGylated proteins through selective autophagy is facilitated by the interaction of ISG15 with p62 and HDAC6. (Nakashima et al., 2015; Villarroya-Beltri et al., 2017). RPS6KA3 (Ribosomal Protein S6 Kinase A3) was identified as an aggresome component as a VIM-APEX2 hit, that has been implicated in the activation of the IRE1 signaling pathway, which in turn regulates HDAC6 activity promoting aggresome formation (Rashid et al., 2015). We also identified the involvement of several proteins that proximally interact in response to stress like CSRP1, LAMP2, DUSP1, SMARCA2, FUS, BAG3, BANF1, SFN, A2M, ISG15, HMGB3, ABCD1, PDLIM1, PPP6C, PDCD4, SMAD2, SNW1, VAMP7, G6PD and RPS6KA3 as aggresome components. Proteins involved in autophagy like LAMP2 (Fang et al., 2022), BAG3 (Gamerdinger et al., 2009) and ATP6V1G1 (Formica et al., 2021) were identified as aggresome components in our dataset. We observed enrichment of SCARB2, known to facilitate the fusion of lysosome with autophagosome (Gleich et al., 2013), as an aggresome component. These data indicate that autophagy regulators are recruited already to aggresomes; this may be functionally relevant to the subsequent aggresome clearance at a later stage during APP.

**Figure 3:**
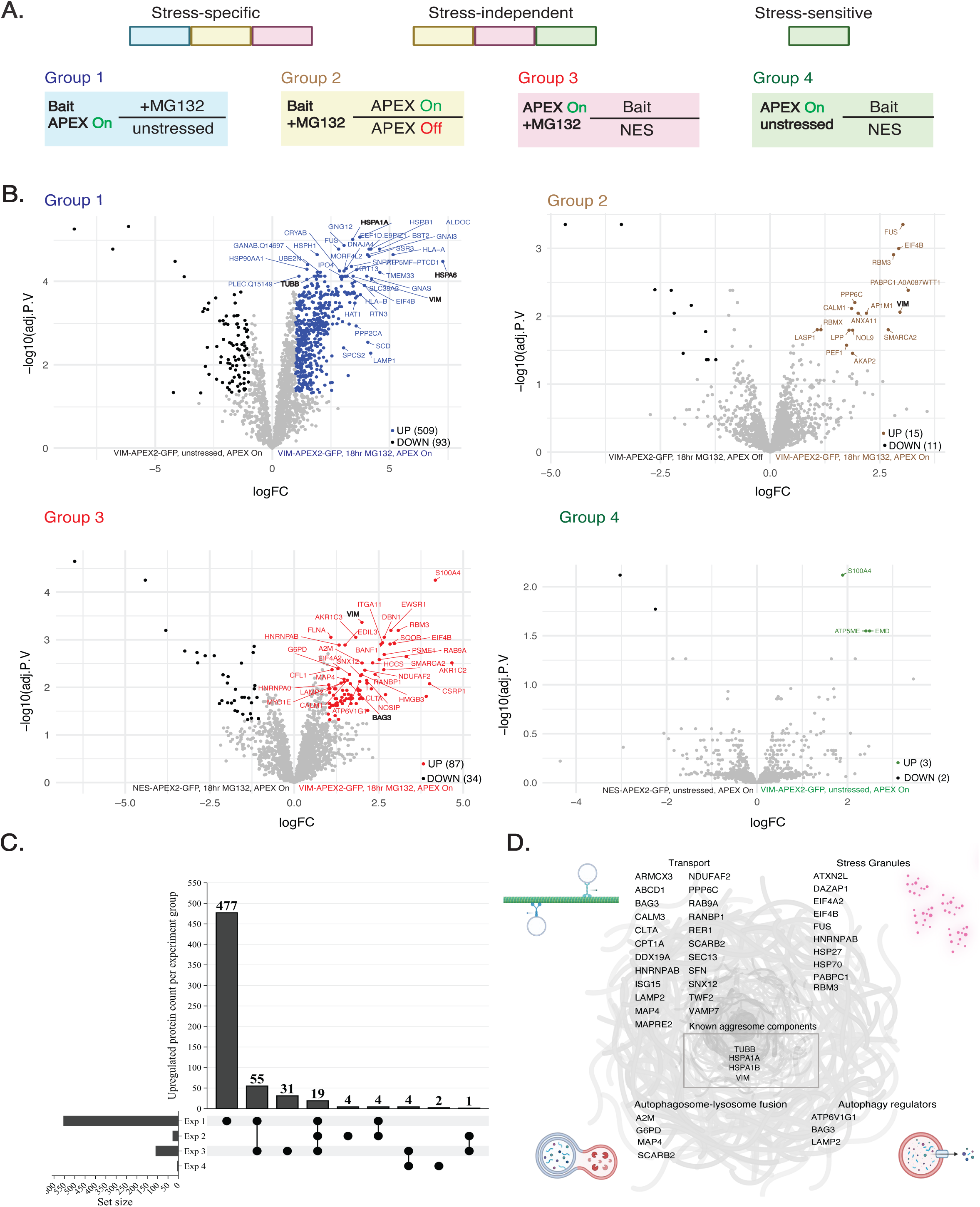
VIM-APEX2 identifies previously known and novel aggresome proteins A) Schematic of comparison groups to detect proteins labelled by the bait under various conditions. The schematic explains each experiment type and the comparisons analysed using the proximity labeling mass spectrometry data set in this study. Further, the schematic describes the classification of 3 types of bait interactors: stress-specific, stress-independent and stress-sensitive, as defined by addition of comparison groups. B) Volcano plots corresponding to each comparison group showing statistically significant (adjusted p value < 0.05) enrichment of proteins in each condition. VIM-APEX2-GFP and NES-APEX2-GFP cells were unstressed or treated with MG132 for 18hr followed by APEX2 mediated biotinylation and streptavidin enrichment of proteins analysed by LC-MS/MS. The top statistically significant upregulated proteins were labelled in each volcano plot and labelled in a colour corresponding to the colour of each experimental group in the schematic in *panel A*. C) Upset plot depicting unique proteins enriched as VIM-APEX2 hits at peak aggresome time point (18hr MG132 treatment) in each comparison group and overlap of shared proteins enriched across 2 or 3 comparison groups. D) Schematic representation of known aggresome proteins and four major classes of novel aggresome proteins identified in our dataset.

### Proximity labeling reveals time-dependent bait interaction networks involved in APP

Next, we performed time-course proximity labeling experiments to determine the changes in molecular interactions during aggresome biogenesis and clearance, induced by application or removal of MG132 stress (Fig.4A). At each time point, we compared the proteins enriched in analysis group 3, as described in Fig.3A, proteins enriched as VIM-APEX2 hits versus the NES-APEX2 background, depicted in the volcano plots (Fig.4B-E). APEX2 samples in triplicates for each time point group clustered separately on a PCA analysis validating our experimental strategy of capturing changes in APP interactome over time (Fig.S6A). At 8 hours of MG132 treatment (early aggresome formation timepoint), we observed an enrichment of proteins like AKR1C3, PPP6C and DHFR in the vicinity of VIM (Fig.S6B). Our analysis also captured DNAJC19 involved in protein folding as a VIM interactor.

**Figure 4:**
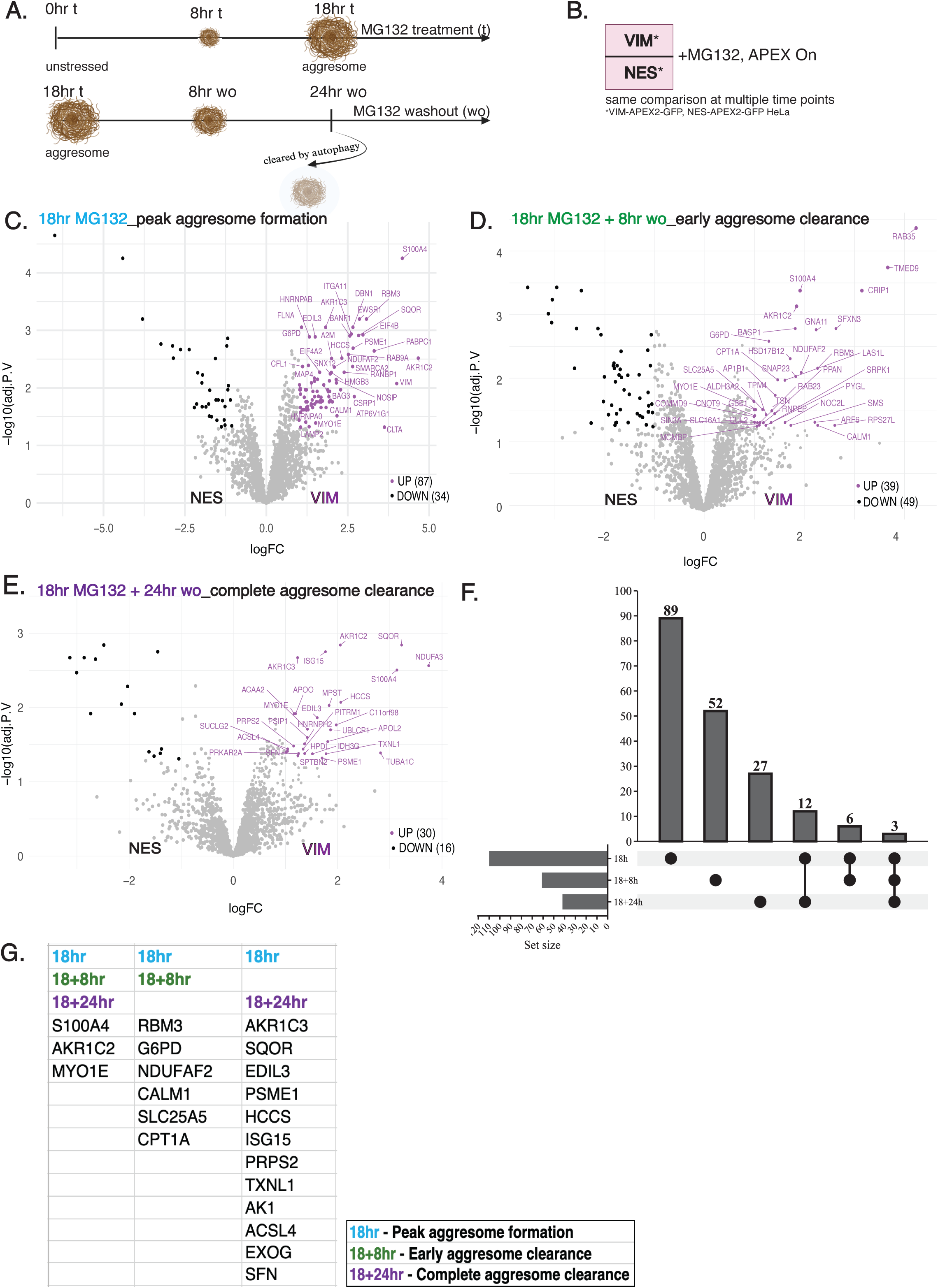
VIM-APEX2 reveals unique and shared proteins during aggresome biogenesis and clearance A) Schematic of time course analysis of APP. The schematic is a representation of the various MG132 treatment and MG132 washout time points that were used to treat VIM-APEX2-GFP, HDAC6-APEX2-GFP and NES-APEX2-GFP cell lines for analysis by proximity labeling mass spectrometry and characterisation of bait protein specific interactome changes in APP. B) Schematic of the analysis groups and the comparison at each time point of APP by proximity labelling mass spectrometry. C-E) Volcano plots showing statistically significant (adjusted p value < 0.05) enrichment of newly identified VIM interactors at different time points during aggresome biogenesis and aggresome clearance. VIM-APEX2-GFP and NES-APEX2-GFP cells were subjected to MG132 treatments and washout as described in *schematic 4A* followed by APEX2 mediated biotinylation and streptavidin enrichment of proteins and analysis by LC-MS/MS. The statistically significant upregulated proteins were labelled in each volcano plot. F) Upset plots showing unique and shared proteins enriched as VIM-APEX2 hits during peak aggresome formation time point and at early and late time points of aggresome clearance. G) List of proteins shared by peak aggresome formation and aggresome clearance time points.

Next, we analysed the proteins detected at “early time point of aggresome clearance”, that is at 8 hours of MG132 washout after mature aggresome has formed. Besides capturing 52 unique proteins at this time point, we also identified 6 proteins like RBM3, G6PD, NDUFAF2, CALM1, SLC25A5 and CPT1A that are components of mature aggresome and were identified during this early aggresome clearance hinting to their potential function in autophagy mediated aggresome clearance (Cho et al., 2025; Giles et al., 2022; Mele et al., 2019; Nakamura et al., 2022; Wang et al., 2023) (Fig.4F-G). GO analysis of molecular function of the enriched unique proteins revealed proteins involved majorly in small molecule binding, hydrolase activity, transferase activity, transporter activity amongst others (Fig.S6C). Several of these 52 unique proteins are involved in establishment of localisation, cellular biological quality, response to stress and autophagy (Fig.S6D). We identified the enrichment of RAB23, CALM1, RAB8A, SLC25A5 and PRKACA as the new autophagy regulators coming in proximity of VIM, particularly at this time point (Fig.S6D, row highlighted in bold black). Interestingly, we observed enrichment of several Rab GTPases in our dataset during the 8 hour washout time point, like RAB35, RAB8A, RAB23 and RAB32 known to be involved in maintaining actin dynamics and intracellular transport and RAB32 is a positive regulator of autophagy (Ao et al., 2014; Hirota & Tanaka, 2009). VIM-APEX2 also identified calmodulin CALM1 and CALM3 which has multi-faceted role in autophagy (Fig.S6D). Specifically, calcium signaling through calmodulins can activate the CaMKKβ-CaMKII pathway, which regulates the activation of ULK1, initiating autophagy under stress conditions (Kim et al., 2011). We used the same approach to identify VIM interactors once the aggresome is completely cleared, that is, at 24 hours of MG132 washout. We identified 27 unique proteins as VIM-interactors when the aggresome is cleared (Fig.4F). Interestingly, there were 12 proteins that followed as VIM interactors from when the aggresome was formed to when it was completely cleared from circulation like AKR1C3, SQOR, EDIL3, PSME1, HCCS, ISG15, PRPS2, TXNL1, AK1, ACSL4, EXOG, SFN. Of these, AKR1C3, EDIL3, ISG15 and ACSL4 have been reported to regulate VIM expression in the context of cellular stress like cancer (Berr et al., 2023; Gasca et al., 2020; Lin et al., 2024; Wang et al., 2025) (Fig.4G). Of note, we did not detect any autophagy regulators when the aggresomes are completely cleared as compared to early time point of aggresome clearance. This explains the specific role of autophagy in initiating the clearance of aggresomes and dissociating from VIM when the aggresomes are cleared.

### Proteasome regulatory proteins are recruited in proximity of HDAC6 and VIM when the aggresome is completely cleared

Aggresomes were cleared from our cell lines after 24 hours of MG132 washout post aggresome formation (Fig.5A). At that time point 42 unique proteins were identified by proximity labeling as HDAC6 interactors, out of which the top 15 are labelled in the volcano plot (Fig.5B). GO analysis of biological function described proteins involved in stress response as top hits like RPA2, TPR, ACAA2, NPM1, ISG15, RFC1, SMC3, ACADM, NIBAN, PSIP1 and TIAL1 (Fig.S7A). Similarly, a GO analysis of molecular function of the 42 enriched proteins revealed proteins majorly involved in unfolded protein binding, dynein binding; however, interestingly the most abundantly enriched proteins are involved in RNA binding such as TIAL1, SRSF11, SRSF7, PABPC1, RBM42, SERBP1, RPA2, TRMT10C, MRPS18B and MRPL2 (Fig.5C). Our dataset also identified myosin, MYO1E and tubulin, TUBA1C in proximal interaction with VIM once the aggresome has been cleared (Fig.S7B).

**Figure 5:**
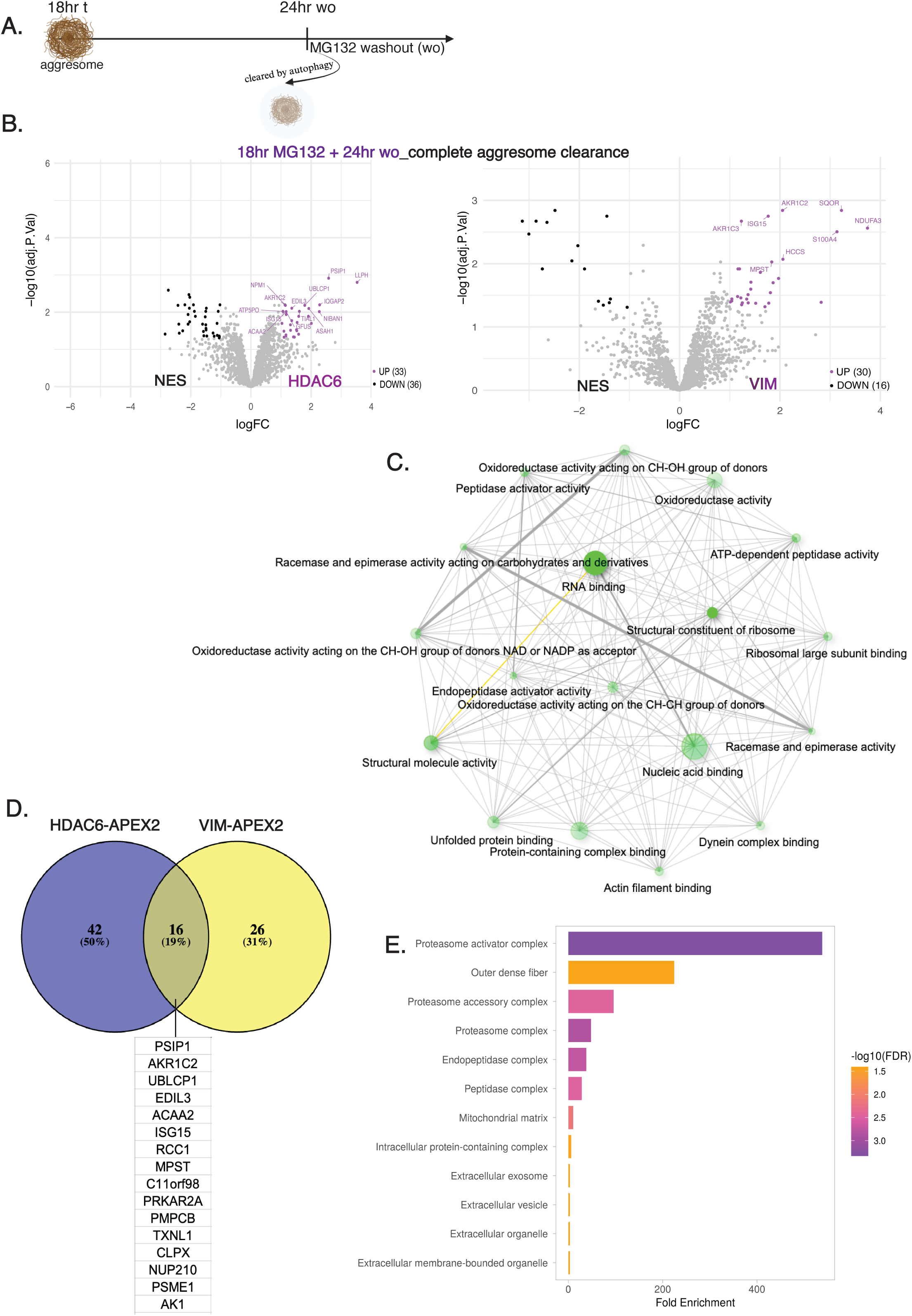
Proximity labeling identifies new HDAC6 and VIM interactors involved in aggresome clearance A) The schematic is a representation of the 18hr MG132 treatment followed by 24hr MG132 washout time point in VIM-APEX2-GFP, HDAC6-APEX2-GFP and NES-APEX2-GFP cell lines for characterisation of bait protein specific interactome changes when the aggresome is cleared. B) Volcano plots showing statistically significant (adjusted p value < 0.05) enrichment of newly identified APP proteins from HDAC6 and VIM point of view when aggresome is cleared. HDAC6-APEX2-GFP, VIM-APEX2-GFP and NES-APEX2-GFP cells were subjected to conditions as described in (A) above, followed by APEX2 mediated biotinylation and streptavidin enrichment of proteins analysed by LC-MS/MS. The statistically significant upregulated proteins were labelled in each volcano plot. C) Dense interaction network of upregulated proteins when the aggresome is cleared and their molecular function. The 42 unique HDAC6-APEX2 hits were used as input in ShinyGO 0.80 software. D) Venn diagram depicting the unique and shared interactors of HDAC6 and VIM during aggresome clearance. The figure was generated using Venny 2.1 software. E) GO analysis of the 16 shared VIM-APEX2 and HDAC6-APEX2 proteins.

A comparison of enriched proteins once the aggresome is completely cleared identified 16 proteins as common interactors of HDAC6 and VIM (Fig.5D). A cellular compartment gene ontology analysis of these 16 shared interactors indicated the maximum enrichment of these proteins as part of proteasome activator complex, proteasome accessory complex and proteasome complex as the top 3 hits (Fig.5E). Amongst the shared HDAC6 and VIM interactors, our dataset identified UBLCP1 (Ubiquitin-like domain-containing C-terminal phosphatase 1) that is known to interact with the 26S proteasome and regulate its activity. UBLCP1 has been shown to dephosphorylate the proteasome, thereby modulating its proteolytic functions (Kors et al., 2019). Another protein that maintains protein quality control in cells by preventing protein aggregation identified in our dataset - ClpX is an ATP-dependent unfoldase chaperone that collaborates with ClpP peptidase to facilitate the degradation of misfolded proteins by unfolding and translocating them into the proteolytic chamber (Maillard et al., 2011). Similarly, PSME1 (Proteasome activator subunit 1), known as PA28α, a regulatory component of the 20S proteasome complex was a shared HDAC6-APEX2 and VIM-APEX2 hit. Its function is to enhance the proteasome’s ability to degrade ubiquitinated proteins, playing a crucial role in maintaining cellular protein homeostasis (Sahu et al., 2021).

## Discussion

Proximity proteomic mapping by multiple baits has been used for studying the composition of membrane-less structures like stress granules (Markmiller et al., 2018; Marmor-Kollet et al., 2020), autophagosomes (Zellner et al., 2021) and lipid droplets (Bersuker et al., 2018). The aggresome is a cluster of ubiquitinated, misfolded aggregated proteins that assembles at the MTOC before being cleared by autophagy. In this study, we have used time-dependent proximity labeling to better understand the dynamics of the APP and to gain insights into the compositional diversity of the mature aggresome. Our study can be used as in-depth resource for the spatiotemporal landscape of interactions during the aggresome processing pathway. Particularly, we provide for the first time an analysis of aggresome formation and clearance at proteome resolution.

We first studied the kinetics of APP by immunofluorescence microscopy. Our data show that small Ub-positive aggregates start to accumulate in HeLa cells at 8 hours of treatment with MG132, a specific inhibitor of the 26S proteasome. We observed that these small aggregates grow in size at 12 hours of MG132 stress and there is a dramatic increase in the number of cells comprising a Ub-positive aggresome at 18 hours of MG132 treatment compared to 12 hours. Therefore, we have used 18 hours of MG132 stress as the peak aggresome time point for our mass spectrometry studies. Aggresomes differ in shapes and size across cell types, and the peak aggresome formation time point here is HeLa specific. The aggresome shape might also slightly differ if visualised using different antibodies-as we also observed when staining aggresomes using the Ub, HDAC6 and VIM antibody, respectively.

Next, we performed dual aggresome bait-VIM and HDAC6 specific proximity labeling, enrichment of biotinylated proteins by streptavidin pulldown and label free quantitative mass spectrometry followed by bioinformatics analysis of the dataset. Already at 8 hours of MG132 treatment, at the early aggresome biogenesis time point, we detected enrichment of PPP6C (protein phosphatase 6 catalytic subunit). It has been reported that the assembly of the WHIP-TRIM14-PPP6C mitochondrial complex promotes RIG-I-mediated antiviral signaling (Tan et al., 2017); this further supports evidence from previous studies that APP and IAV uncoating have shared molecular players (Wang et al., 2022). At 18 hours of MG132 treatment, we discovered more than 75 new proteins in the mature aggresome, establishing a resource of high confidence by comparison among 4 different groups. GO analysis showed 23 of these new aggresome components were involved in protein transport; although further experimental validation is required, we hypothesize that these proteins in interaction with VIM facilitate the transport of aggregated proteins towards the MTOC. We identified disease relevant proteins as VIM interactors in the aggresome. For example, we identified ANXA11 as an aggresome component. Recent studies have highlighted ANXA11’s role in protein homeostasis, particularly in the context of neurodegenerative diseases. Mutations in the ANXA11 gene have been associated with amyotrophic lateral sclerosis (ALS) and certain myopathies (Smith et al., 2017). Another new aggresome component, A2M has been shown to interact with aggregation-prone proteins, facilitating their internalization and degradation via the lysosomal pathway. This function positions A2M as a clearance factor that aids in the removal of potentially toxic protein aggregates from cells (Tomihari et al., 2023). Further, DUSP1 was also identified in our list of aggresome proteins – DUSP1 (Dual-specificity phosphatase 1), also known as mitogen-activated protein kinase phosphatase-1 (MKP-1), is a key regulator of mitogen-activated protein kinase (MAPK) signaling pathways, including JNK, p38, and ERK. By dephosphorylating these kinases, DUSP1 modulates various cellular processes such as inflammation, stress responses, and cell differentiation (Bhore et al., 2017). Another interesting observation was the enrichment of G6PD (Glucose-6-phosphate dehydrogenase) as an aggresome component, which is the rate-limiting enzyme of the pentose phosphate pathway, crucial for cellular defense against oxidative stress through NADPH production. Alternatively, acetylation of G6PD has been linked to its stability and aggregation propensity. Specific acetylation sites on G6PD influence its aggregation over time, which could have implications for cellular stress responses and protein quality control mechanisms (Wu et al., 2023). We identified MAP4 (Microtubule associated protein 4) involved in microtubule stability as an aggresome component, in line with previous studies that reported the role of MAP4 in APP in a double role: MAP4 inhibition leads to aggresome formation and MAP4 localisation at the perinuclear reagion attracts lysosomes to the aggresome site promoting autophagic clearance (Lazaro-Dieguez et al., 2008; Zaarur et al., 2014). Interestingly, RPS6KA3 (Ribosomal Protein S6 Kinase A3) was identified as an aggresome component as a VIM-APEX2 hit, that has been implicated in the activation of the IRE1 signaling pathway, which in turn regulates HDAC6 activity promoting aggresome formation (Rashid et al., 2015).

Our dataset also validates previous evidence stating that rigid stress granules are cleared by autophagy. We identified ATXN2L, DAZAP1, EIF4A2, EIF4B, FUS, HNRNPAB, PABPC1 and RBM3 as mature aggresome components that were also identified as stress granule components (Markmiller et al., 2018; Marmor-Kollet et al., 2020). Additionally, our data is consistent with published evidence that reported “aberrant SGs”, or SGs containing misfolded proteins are positive for chaperones such as Hsp70 and Hsp27, also identified in this study as components of the mature aggresome. The authors reported that these aberrant SGs containing ALS associated-misfolded SOD1 aggregates are cleared by aggresome pathway dependent autophagic degradation (Mateju et al., 2017). These findings indicate a strong association between SGs and APP, highlighting potential avenues for future investigation.

Our analysis revealed many autophagy regulators as new components in the mature aggresome with functions ranging from recruitment of autophagosomes to the MTOC and fusion of lysosomes with autophagosomes that facilitate autophagic clearance. Besides LAMP2, BAG3, ATP6V1G1, we observed upregulation of SCARB2, known to facilitate the fusion of lysosome with autophagosome (Gleich et al., 2013), as an aggresome component. Similarly, we observed upregulation of VAMP7 that is known to mediate lysosome fusion to autophagosome (Jian et al., 2024) and regulate cytoskeleton dynamics (Koseoglu et al., 2015) thus promoting APP. These data indicate that autophagy regulators are recruited early to aggresomes, and a future direction is to validate aggresome localisation of these hits by immunofluorescence microscopy. This engagement of at least some of the proteins responsible for autophagy may be functionally relevant to the regulation of aggresome clearance at a later stage.

In addition, our data illustrates the dynamics of several proteins that accumulate in the aggresome and further drive the aggregated proteins to autophagic clearance. At early aggresome clearance time point, we observed enrichment of PGAM5, a protein involved in autophagy. It has been reported that in response to mitochondrial stress PGAM5 binds and dephosphorylates FUNDC1, promoting recruitment of autophagosomes to mitochondria (Ruiz et al., 2019). At the same time point, GO analysis revealed identified proteins like MTMR2, GNA11, RAB23, RAB32, SAR1B, RNPEP, RAP2B, RAB35, MYO1E, ARF6, RAB8A, LAS1L, AQR, GBE1 and TSN with hydrolase activity and CSNK1A1, GBE1, SRPK1, PYGL, SMS, CPT1A, PRKACA, CPNE3, GTF3C4, CALM3 and CALM1 with transferase activity. We hypothesize that these hydrolases are involved in different stages of the autophagy process, particularly in the degradation of autophagic cargo within the lysosome and that transferases play essential roles in autophagy by facilitating the modification of proteins and lipids required for autophagosome formation, cargo recognition, and lysosomal degradation. We observed enrichment of several Rab GTPases in our dataset at the 8 hour washout time point. Amongst them, RAB32 is a positive regulator of autophagy. Our result validates previous evidence that mutations in RAB32 lead to formation of Ub positive aggregates (Ao et al., 2014; Hirota & Tanaka, 2009). VIM-APEX2 also identified calmodulin CALM1 and CALM3 which has multi-faceted role in autophagy (Fig.D). Specifically, calcium signaling through calmodulins can activate the CaMKKβ-CaMKII pathway, which regulates the activation of ULK1, initiating autophagy under stress conditions (Kim et al., 2011). Additionally, the identification of both CALM1 and CALM3 during early aggresome clearance time point is a positive validation of our proximity labeling dataset as they are known interactors of VIM (Campetelli et al., 2012).

When the aggresomes are completely cleared from cells, we observed the recruitment of common HDAC6 and VIM specific interactors that are components of the proteasome activator and accessory complex like UBLCP1, PSME1 and ClpX. This suggests that upon eliminating cellular toxicity through aggresome clearance, the cell becomes primed for a subsequent cycle of protein degradation.

Future work should focus on identifying the critical factors and mechanisms in APP. The APP protein dataset we present here suggests possible future directions and provides a framework for identifying previously unknown important regulators. We present the example of several autophagy regulators in interaction with VIM, which are not only essential for aggresome formation (LAMP2, ATP6V1G1 and BAG3) but also for aggresome clearance (RAB23, CALM1, RAB8A, SLC25A5 and PRKACA). These can be promising therapeutic targets and strategies will therefore likely need to specifically target those mechanisms in disease relevant cell lines. Our work represents a step along this path, which so far has been hindered by a sparsity of promising targets. Future studies include validation of identified hits by – first, validating aggresome localisation by immunofluorescence microscopy and second, siRNA knockdown of identified hits to check if the APP is hindered. Additionally, by using a combination of different APEX2-tagged validated hits from this study will make it possible to further dissect the molecular architecture of APP and enable the distinction as well as the characterization of cell-type-specific aggresomes such as in neurodegenerative disease models or in models to study RNA virus uncoating.

## Acknowledgements

We thank Laure Plantard and Laurent Gelman for setting up and helping with the microscopy experiments. We thank Hubertus Kohler for help with FACS cell sorting. We thank Charlotte Soneson for help with the R script for generation of volcano plots. The schematics were created with BioRender.com.

## Authors contributions

Conceptualization: SG, LW, PM Methodology: SG, LW, DH, VL, JS, JE, CC Investigation: SG, DH, JS, JE Visualisation: SG, DH, JS

Funding acquisition: PM Project administration: GM, PM Supervision: PM

Writing – original draft: SG, PM Writing – editing: SG, PM

## Data and materials availability

All unique reagents generated in this study are available from the lead contact with a completed Materials Transfer Agreement (MTA); published research reagents from the FMI are shared with the academic community under an MTA having terms and conditions corresponding to those of the UBMTA (Uniform Biological Material Transfer Agreement). Microscopy data reported in this paper will be shared by the lead contact upon request. Any additional information required to reanalyse the data reported in this paper is available from the lead contact upon request.

## Supplementary Materials

Materials and Methods

Figs S1-S7

