## Supplementary data for "Dynamic profiling of the aggresome processing pathway"

**Affiliations:**

**The PDF file includes:**

Materials and Methods

Figs. S1 to S7

#### Material and Methods

##### Cell lines and culture conditions

HeLa and HEK293T cells were grown and cultured in DMEM medium supplemented with 10% fetal bovine serum. All cells were cultured in a humidified incubator at 37°C with 5% $CO_2$. All cell lines were routinely tested for mycoplasma.

##### Construct design and cloning of constructs

All pcDNA 3.1 constructs were cloned using NEB Hifi DNA Assembly kit (NEB Catalog #E2621S). The protocol used was as follows- The FLAG-HA-pcDNA3.1 plasmid backbone (Addgene plasmid #52535) was double-digested at the restriction enzyme sites Xba1 (NEB Catalog #R0145S) and BamH1 (NEB Catalog #R3136S) using the restriction-digestion protocol from NEB and 10X NEB buffers (Catalog #B7030). The inserts were PCR amplified from respective plasmid backbones- hsHDAC6 from pOPINF-HIS6-3C-hsHDAC6, VIM from EGFP-Vimentin-7 (Addgene plasmid #56439), both APEX2 and copGFP fragments from HR_G3BP1-V5-APEX2-GFP (Addgene plasmid #105284) and mGreenLantern from pcDNA3.1-mGreenLantern (Addgene plasmid #161912). The PCR amplified template DNA and the double-digested plasmid DNA were run on 1% agarose gel and DNA bands of the correct size was extracted using the gel extraction kit following the manufacturer's protocol (Promega; Catalog #A9281). The Gibson assembly reactions were set up according to the NEB Hifi DNA Assembly kit manufacturer's protocol. The lentivirus constructs are from Vector Builder- HDAC6-APEX2-GFP (Vector name:pLV-Puro-hPGK>(R-Kozak-Flag-HDAC6-APEX2-mGL-R)/3xGS,Catalog#: Ecoli(VB230417-1234uag)) , VIM-APEX2-GFP (Vector Name: pLV-Puro-hPGK-xba1-flag-vim-apex2-mgl-sal1, Catalog #:Ecoli(VB230815-1215sxv)) and NES-APEX2-GFP (Vector name: pLV-Puro-hPGK>(xba1-flag-APEX2-NES-mGL-sal1), Catalog #:Ecoli(VB230816-1213zrc)).

##### Transfection

HEK293T cells and HeLa cells were transfected with FuGENE HD (Promega, Catalog #E2311) following the manufacturer's protocol.

##### CRISPR KO cell line generation

To generate HDAC6 CRISPR KO clones, wild-type HeLa cells (ATCC) were transfected with pX330-Cas9-T2A-mCherry expressing the sgRNA 5'-GGTGGAATCCTGGCCGGTTG-3'. targeting a sequence within exon 2 of hsHDAC6, enhancing efficiency of KO cell line generation (Saito et al., 2019). Transient transfections were performed following the manufacturer's instructions using FuGene HD and DMEM medium supplemented with 10%

fetal bovine serum (Thermo Fisher Scientific). 48 hours post transient transfections, cells expressing high mCherry level were pool sorted for western blot analysis for sgRNA effciency testing and single-cell–FACS sorted into 96-well plates for monoclonal selection (10% highest mCherry-positive cells, using Cas9-T2A-mCherry). Single clonal cell populations were expanded over weeks and screened for loss of HDAC6 by immunostaining with HDAC6 antibody (HDAC6 (D2E5) Rabbit mAb CST#7558) and acetylated alpha-tubulin antibody (Sigma, Clone 6-11B-1 #T745).

To generate VIM CRISPR KO clones, wild-type HeLa cells were similarly transfected with pX330-Cas9-T2A-mCherry expressing the sgRNA 5'-GCCGAGCTCGAGCAGCTCAA-3'. Transient transfections were performed following the manufacturer's instructions using FuGene HD and DMEM medium supplemented with 10% fetal bovine serum (Thermo Fisher Scientific). 48 hours after transient transfections, highly transfected cells were pool sorted for western blot analysis for sgRNA effciency testing and single-cell–FACS sorted into 96-well plates for monoclonal selection (10% highest mCherry-positive cells, using Cas9-T2A-mCherry). Single clonal cell populations were expanded over weeks and screened for loss of VIM by immunostaining with VIM antibody (Vimentin (D21H3) XP® Rabbit mAb CST#5741).

For both HDAC6 and VIM KO cell lines, several KO clones each were verified by western blot for depletion of respective protein expression and by genotyping PCR.

**Generation of APEX2 cell lines**

Transfer plasmid pLV-Puro-hPGK-Flag-HDAC6-APEX2-GFP or pLV-Puro-hPGK-Flag-VIM-APEX2-GFP or pLV-Puro-hPGK-Flag-NES-APEX2-GFP were co-transfected with packaging plasmids (expressing tat, rev, gag, vsv-g) together with FuGENE HD reagent (Promega#E2311) for establishment of respective HeLa cell lines. After 25 min incubation at room temperature, the mix was added to HEK293T cells in 10 cm culture dish, seeded one day before with 1.6 million cells per dish. Cells were cultured at 37°C for 3 days; the medium was collected and filtered with 0.45 mm filter (Merck#SEIM003M00), and LentiX concentrator (Takara#631231) was added (1/3 of the supernatant volume). The mixture was incubated at 4°C for 30 min and the lentivirus was precipitated by centrifugation at 1500 g, 45 min, 4°C. The pellet was re-suspended in 500mL Opti-MEM (Gibco#31985062). The re-suspended lentivirus pellet was added to the culture medium of HDAC6 KO or VIM KO HeLa cells (10 cm dish, 1.6 million cells seeded per dish one day before). GFP positive cells were then single cell sorted into 96 wells plates. After 1 month of culturing, clones were expanded and analyzed by western blot with respective antibodies: HDAC6 antibody (HDAC6 (D2E5) Rabbit mAb

CST#7558), VIM antibody (Vimentin (D21H3) XP® Rabbit mAb CST#5741) and GFP antibody (GFP Polyclonal Antibody Invitrogen#A-11122).

**Aggresome induction and clearance in cell lines**

10μm MG132 (Sigma Aldrich Cat# M7449) supplemented in complete DMEM medium was used to treat cells for 18 hours to induce aggresome formation for both immunofluorescence and proximity labeling experiments. To study aggresome clearance, cells were first treated with MG132 to induce aggresome formation followed by washing the cells with complete DMEM medium several times and then maintaining the cells in MG132 free complete medium for 24 hours in a humidified incubator at 37°C with 5% $CO_2$.

**Western blot**

Treated HeLa cells were lysed with RIPA buffer (supplied with cOmplete Protease Inhibitor Cocktail, Merck#11697498001), and protein concentration was determined by Bradford assay. The same protein amounts were loaded onto 4 to 12% NuPAGE Gels and protein was transferred to PVDF membranes by using Trans-Blot Turbo Transfer System (Biorad#17001917). The membrane was blocked with 5% BSA in TBS and then incubated with primary antibodies overnight at 4°C. The signal was developed with HRP-conjugated secondary antibodies and Amersham ECL Western Blotting Detection Reagent (Cytiva#RPN2106). Antibody information: HDAC6 antibody (HDAC6 (D2E5) Rabbit mAb CST#7558; 1:1000 dilution), VIM antibody (Vimentin (D21H3) XP® Rabbit mAb CST#5741; 1:1000 dilution) and GFP antibody (GFP Polyclonal Antibody Invitrogen#A-11122; 1:1000 diltuion), Streptavidin, horseradish peroxidase conjugate (Invitrogen, Cat# S911 (Lot Number 2384052); 1:2000 dilution), alpha-Tubulin antibody (Rabbit polyclonal, Abcam#18251; 1:1000 dilution), Acetylated alpha-Tubulin antibody (Sigma, Clone 6-11B-1 #T745; 1:2000 dilution), Actin antibody (CST#4967; 1:2000 dilution), ECL anti-rabbit IgG, HRP linked whole antibody (from donkey, Lot: 17457635 (NA934V); 1:5000 dilution), ECL anti-mouse IgG, HRP linked whole antibody (from sheep, Lot: 17479274 (NA931V); 1:5000 dilution).

**APEX2-mediated biotinylation and enrichment of biotinylated proteins**

Hela cells (APEX2-GFP cell lines) were seeded in 6cm culture dishes one day prior to be 80% confluent the following day. Cells were treated with 10uM MG132 or left unstressed and incubated for the duration of stress in a humidified incubator at 37°C with 5% CO2. For proximity labeling, the culture medium was aspirated, and cells were replaced with complete DMEM medium supplemented with 2.5mM biotin-phenol (BP) for 30min except for the no-substrate control samples. Cells were washed with phosphate buffer saline (PBS) 3 times.

APEX labeling was performed by adding hydrogen peroxide ($H_2O_2$) to a final concentration of 1mM for 2min before quenching the biotinylation reaction by adding quenching buffer containing Trolox ((+/ )-6-Hydroxy-2,5,7,8-tetramethylchromane-2-carboxylic acid, Sigma Cat#238813) to a final concentration of 5mM and sodium L-ascorbate (Sigma Cat#A4034) to a final concentration of 10mM dissolved PBS. Cells were washed twice with cold PBS and collected using cell scrapers and pelleted for 3min at 300g. Immediately after, the cells were lysed by suspending in 1mL ice cold lysis buffer containing 8M urea, 150mM NaCl, 20mM Tris-HCl pH 8.0, Protease Inhibitor Cocktail Set III, EDTA-Free (EMD Millipore, Cat#539134), 5mM Trolox and 10mM sodium L-ascorbate. The samples were sonicated (5sec On, 5 sec Off – 3 times, for each tube) and cleared by centrifugation at 12000rpm for 10min at 4°C. The protein was transferred to new 1.5mL tubes. Protein concentration was measured by Bradford Assay. For affinity purification, we used 10uL of streptavidin magnetic beads for 200ug protein (Pierce, Cat#PI88817). Beads were once in 2M urea buffer. 200ug protein was incubated with the beads. The samples were diluted to 2M urea by adding 3 volumes of 150mM NaCl, 20mM TrisHCl pH 8.0 with protease inhibitors and quenchers. Samples were incubated on a rotating stand for 2hr at room temperature and washed 8 times in 2M urea buffer followed by 3 times with Tris-buffered saline (TBS). Following the washes, the beads at 240 RCF for 5 min at 4°C Can check now for correct and effective biotinylation by WB (for WB, elute proteins from beads by boiling in loading dye supplemented with biotin, run gel, check by Strep-HRP antibody).

**Mass spectrometry sample preparation- Reduction, alkylation and digestion**

20uL of Lysine-C stock was added to 100uL of digestion buffer. The wash buffer was completely from the beads by pipetting and 6uL of Lys-C in digestion buffer was added, the samples were spinned down quickly (<1000xg) and incubated at room temperature for 2-4 hours while shaking, then spinned down. 20uL of 0.2ug/uL trypsin was mixed with 340uL 50 mM HEPES and 18uL of the trypsin mix was added to beads, spinned down, vortexed, spinned down again and incubated overnight at 37°C in an incubator. The next morning, samples were spinned down, 1uL 0.2ug/uL trypsin was added, spinned down, vortexed, spinned down again and the digestion step was continued for additional 4-6 hours in an incubator. After digestion, samples were spinned down and processed immediately or frozen at -80°C to be analyzed by MS later (pause point).

**Mass spectrometry LC-MS/MS**

Vanquish Neo with 2 column setup, 15cm EASY Spray

"D:\Data\LUMOS\Methods\Tune4.0_IP_L3\IP_80min_200nl_NTC_HCD_IT_230530.meth"

Peptides generated by Lys-C/trypsin digestion were acidified with 0.8% TFA (final) and analyzed by LC–MS/MS on a Vanquish Neo chromatography system with a two-column set-up (0.3x5 mm Pepmap C18 trapping column and an EASY-Spray Pepmap Neo C18 2um 75umx150mm column heated to 45°C) mounted on an EASY-Spray™ source connected to an Orbitrap Eclipse mass spectrometer (all Thermo Scientific). The peptides were loaded onto the trapping column in 0.1% formic acid and 2% acetonitrile in $H_2O$ at a constant flow rate of 120 µl/min at max. 800 bar. Using a flow rate of 200 nl/min, peptides were then separated at room temperature over a 80 min gradient with a mobile phase buffer system consisting of solvent A: 0.1% formic acid; solvent B: 0.1% formic acid in 80% acetonitrile in a linear gradient of 2–6% solvent B in 1 min followed by a linear increase from 6 to 20% in 42 min, 20–35% in 22.5 min, 35–50% in 3 min, 50-100% in 1.5 min, and the column was finally washed for 10 min at 100% solvent B.
MS1 survey scans were performed every 3s using a 120k resolution (at 200 m/z) in the Orbitrap from 375-1575 m/z in profile mode, with a maximum injection time of 50 ms. Precursors were selected for MS2 based on advanced peak determination, monoisotopic peak determination set to peptides, within charge states 2-7, dynamic exclusion of 8 s within +/-10 ppm tolerance for isotopes, with a minimum intensity of 5e3. Precursors were selected in the quadrupole within an isolation window of 1.6 m/z for MS2 HCD fragmentation with normalized collision energy 30% dynamically setting the max. injection time in rapid mode, in the “normal” mass range in centroid mode.

**Primary data analysis**

FragPipe was chosen as the analysis tool here because of its speed compared to MaxQuant, results are highly comparable. Duration: ALL JOBS DONE IN 328.1 MINUTES
W:\scratch\ganaly_proteomics\gmatthia\ghossuch_793_4551\JS\FragP

**Database search and label-free (LFQ) quantification with FragPipe**

MS raw data were analyzed with FragPipe (https://fragpipe.nesvilab.org/ ) version 22.0. MS2 fragment ion spectra were searched with MSFragger (Kong et al., 2017; Teo et al., 2021) version 4.1 against a protein database consisting of a concatenation of the HUMAN Uniprot proteome (UP000005640_9606, 81,791 entries, downloaded on Jan 12 2023) and a contaminant database (contaminant tag “contam_”), and their reversed decoy complement (decoy tag “rev_”), using default settings: peptide length 7-50 amino acids, peptide mass 500-5000 Da and strict trypsin cleavage specificity [KR, C-terminal, 2 missed cleavages], fixed cysteine carbamidomethylation (C, 57.994), variable methionine oxidation (M, 15.9949) and protein N-terminal acetylation (N-term, 42.0106). Precursor mass tolerance was set to +/-20 ppm, and fragment mass tolerance to 0.6 Da (250 ppm after optimization). PSMs were validated with Percolator (Kall et al., 2007), MS1 label-free quantification was performed with IonQuant (Yu et al., 2021) version 1.10.27 to calculate MaxLFQ intensity based on a min. of 2 ions, and Top3 intensity based on the three most abundant ions, without match-between runs (MBR), including normalization. Protein inference was achieved with ProteinProphet (Nesvizhskii et al., 2003) and FDR filtering and reporting with Philosopher (da Veiga Leprevost et al., 2020).

**Secondary data analysis**

Einprot data analysis (FragPipe LFQ APMS workflow)

"W:\scratch\ganaly_proteomics\gmatthia\ghossuch_793_4551\JS\einprot_4551\reRun\FragPipe_4551_MaxLFQ.intensity_MinProbGlobal_center.median_limma_adjpval_0.1_einprot0.9.6.html"

**Protein filtering, abundance log transformation, normalization, imputation and PCA**

The FragPipe proteins feature table file Proteins.txt was imported into the in-house developed einprot R package version 0.9.6 (Soneson et al., (2023). einprot: flexible, easy-to-use, reproducible workflows for statistical analysis of quantitative proteomics data. Journal of Open Source Software, 8(89), 5750, https://doi.org/10.21105/joss.05750), removing proteins identified from the “Contaminants” database, reverse hits, and proteins only identified by site, and proteins identified with a score lower than 10 or less than 2 peptides. The MaxLFQ.Intensities were log2-transformed, and missing values were imputed using a modified version of the MinProb algorithm implemented in the imputeLCMD R package (https://cran.r-project.org/web/packages/imputeLCMD/index.html), sampling values to impute from a normal distribution with parameters derived from all observed values (MinProbGlobal), rather than separately for each sample. Principal component analysis was performed on the imputed log-transformed intensities using the scater R package (McCarthy et al., 2017).

**Statistical testing using Limma with sample comparisons to broad sample complement or selected sample control groups**

The limma R package (Ritchie et al., 2015) was used to fit a linear model for each group, comparing the imputed log2-transformed intensities from different selected sample groups. For each comparison, at least two observed (non-imputed) values were required to report a test result for a protein. An adjusted p-value threshold of 0.1 was chosen to call significant features in all comparisons.

**Quality check**

Individual samples for 8h_VIM (S33 and S34, 0n and 0ff) did not cluster with the rest of their groups. In an einprot re-run they were excluded from the samples.

**Immunofluorescence and Microscopy**

0.05 million cells were seeded in a 4-well chamber slide (ThermoFisher Cat#154526). Following overnight culture, cells were treated with MG132 for different time points. Subsequently, the medium was aspirated, and cells were washed twice with ice-cold PBS. Cells were then fixed with 4% paraformaldehyde (Ready-to-Use Fixative, Biotium Cat#22023) for 10 min, permeabilized with 0.5% Triton X-100/PBS for 15 min, and blocked with blocking buffer: 10% Normal Goat Serum blocking solution (Invitrogen Cat# 50197Z) for 1 hr at room temperature. Primary antibodies were applied to cells at a 1:100 dilution and incubated overnight at 4°C. The next day, cells were washed three times with PBS and incubated with fluorescent secondary antibodies in blocking buffer (1:100 dilution) for 1hr at room temperature. DAPI (ThermoFisher Cat#D1306) and Actin staining with Phalloidin 488 (CytoSkeleton Cat#PHDG1) were diluted in PBS according to the product instructions and added to cells for 20 min at room temperature. Following three washes with PBS, cells were sealed with ProLong™ Diamond Antifade Mountant (ThermoFisher Cat#P36965). Slides were inspected with a spinning disk confocal microscope unit Yokogawa CSU W1 with Dual T2.

**Image analysis**

Image processing code for the MG132 time-course data has been deposited at Zenodo and is publicly available at https://doi.org/10.5281/zenodo.16845480.

Cell nuclei were segmented on the DAPI channel using Cellpose (Stringer et al., 2021) with the 'nuclei' pre-trained model (parameters: diameter=150, flow_threshold=0.8, cellprob_threshold=0.0). A cell mask was created by expanding the nuclei labels (distance=50). Subsequently, Ub-positiive aggregates were measured in the Ub channel by using a threshold derived from the image mean intensity + 4.5 times the intensity standard deviation. Aggregates with an area>1000 pixels were classified as aggresomes, and the ratio of aggresome-containing cells was calculated as the number ratio of aggresomes and cells (n_aggresomes / n_cells) within each image.

#### Supplementary figures

### Fig.S1.

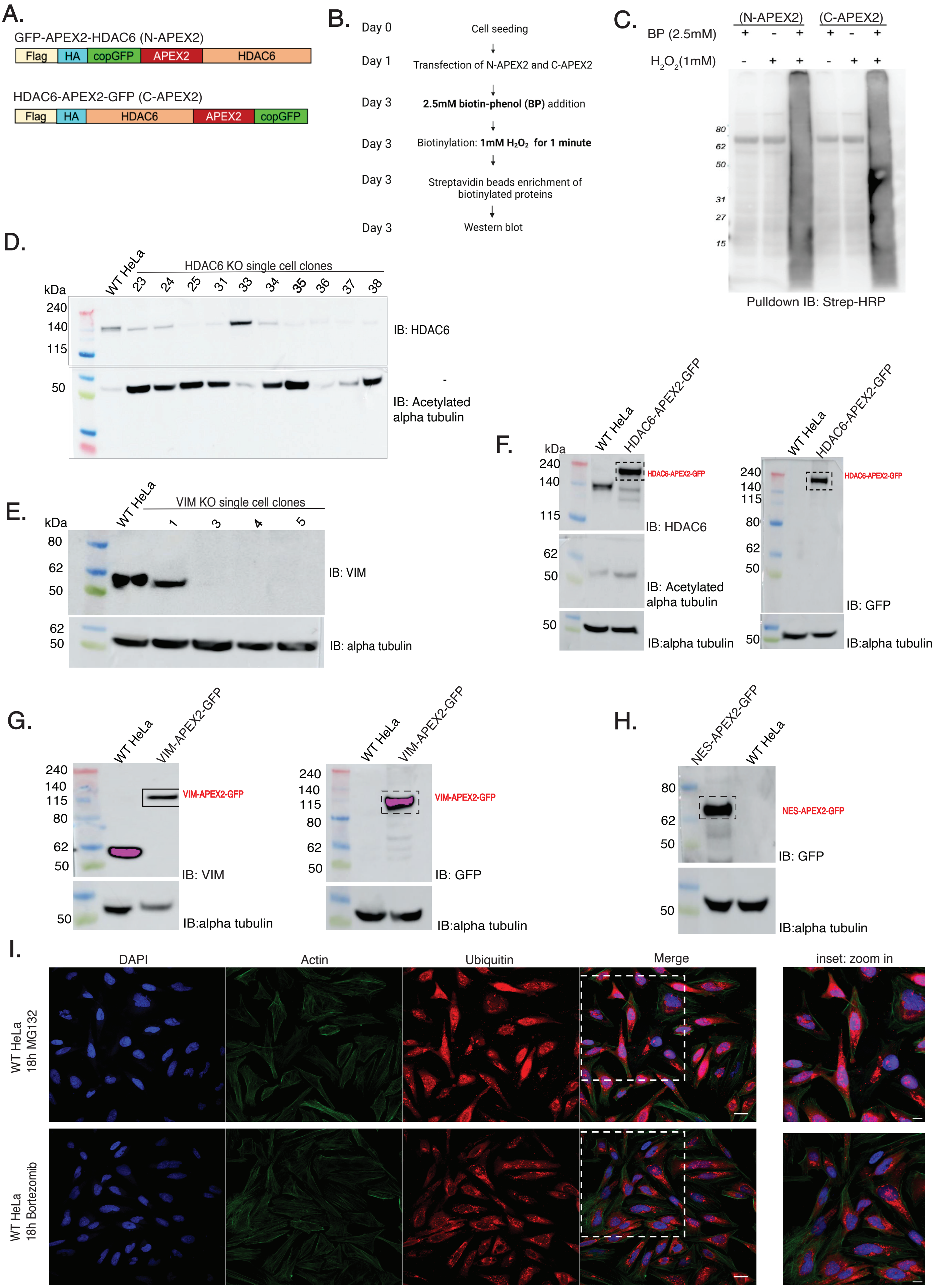

**Figure S1: Testing the functionality of APEX2-copGFP fusion to hsHDAC6, validation of HDAC6 KO and VIM KO cell lines, fusion protein expression from aggresome bait-APEX2-GFP cell lines and MG132 treatment time points for aggresome visualisation in HeLa cells** *(related to Figure 1).*

A) Construct design of N and C terminal APEX-copGFP fusions to hsHDAC6.

B) Schematic of sample preparation to test to test APEX2 activity of above constructs.

C) Immunoblot to confirm functional APEX2 activity of fusion constructs. The smear corresponds to biotinylated proteins pulled down by streptavidin beads and is only visible in the presence of both substrates of APEX2 (BP and hydrogen peroxide) confirming APEX2 activity of the fusion constructs transfected in HDAC6 KO HEK293T cells.

D) Immunoblot to confirm HDAC6 KO single cell clones. Absence of HDAC6 protein and enhanced levels of acetylated of α-tubulin were detected in HDAC6 KO clones compared to WT HeLa. Clone 35 was used for introduction of the HDAC6-APEX2-GFP construct.

E) Immunoblot to confirm VIM KO single cell clones. Absence of VIM protein was detected in VIM KO clones, confirming knockout of VIM in HeLa. Clone 4 was used for introduction of the VIM-APEX2-GFP construct.

F) Immunoblot to test fusion protein HDAC6-APEX2-GFP expression and deacetylase activity of single cell clone H3, by immunobloting with HDAC6, acetylated alpha-tubulin and GFP antibody. This clone was selected for further experiments from multiple clones that were examined similarly.

G) Immunoblot to test fusion protein VIM-APEX2-GFP expression of single cell clone M3, by immunobloting with VIM and GFP antibody. This clone was selected for further experiments from multiple clones that were examined similarly.

H) Immunoblot to test fusion protein APEX2-NES-GFP expression of single cell clone H1, by immunobloting with GFP antibody. This clone was selected for further experiments from multiple clones that were examined similarly.

I) DAPI, Actin and Ubiqutin antibody staining of WT HeLa cells treated with proteasome inhibitors, either 10uM MG132 for 18 hours or 5uM Bortezomib for 18 hours. Ub-positive aggresomes were visualised in both conditions, validating that 18 hours of cellular stress is optimum time point to visualise aggresomes in HeLa cells.

Fig.S2.

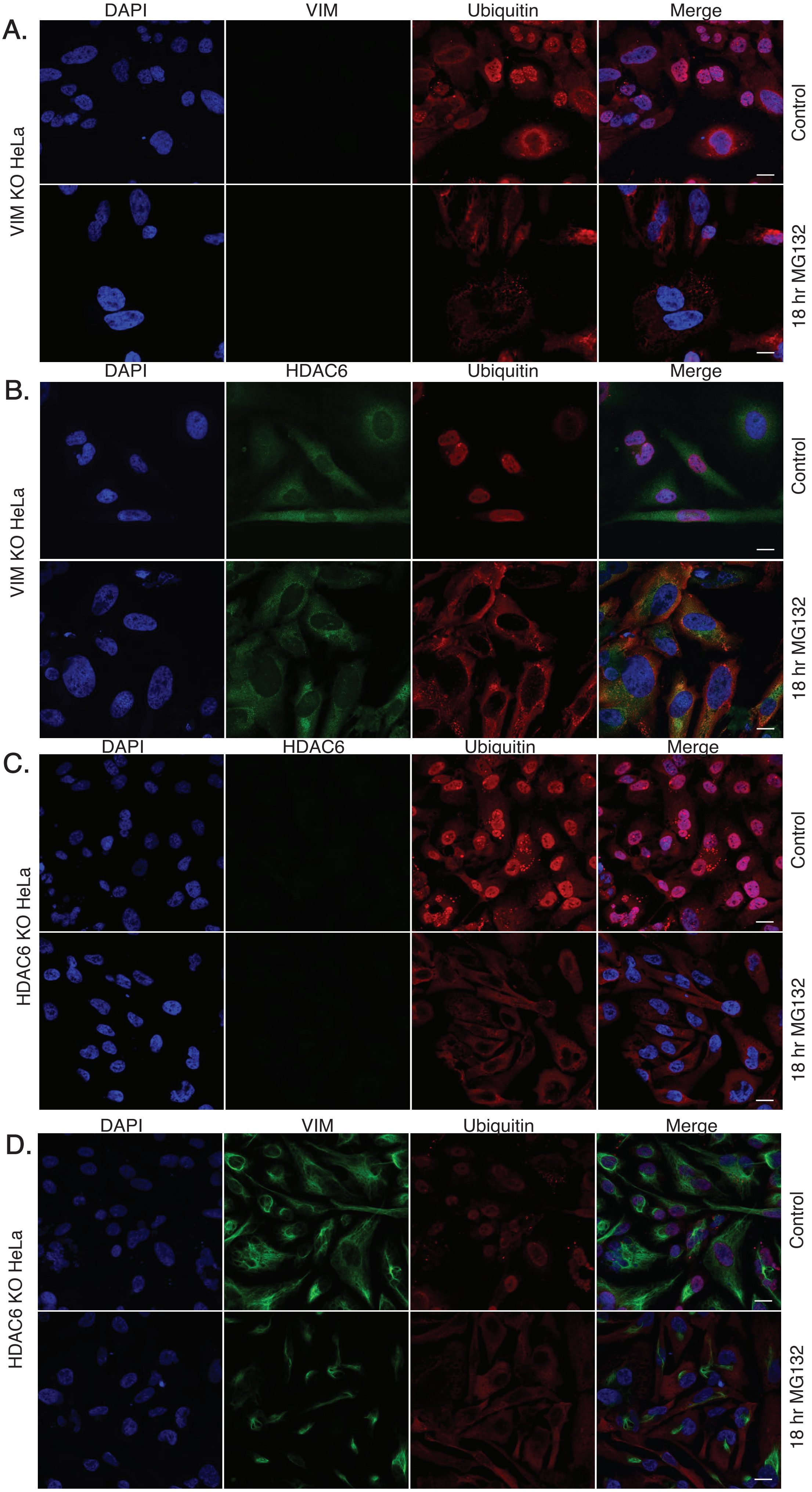
A.
DAPI
VIM
Ubiquitin
Merge
VIM KO HeLa
Control
18 hr MG132
B.
DAPI
HDAC6
Ubiquitin
Merge
VIM KO HeLa
Control
18 hr MG132
C.
DAPI
HDAC6
Ubiquitin
Merge
HDAC6 KO HeLa
Control
18 hr MG132
D.
DAPI
VIM
Ubiquitin
Merge
HDAC6 KO HeLa
Control
18 hr MG132

**Figure S2: Full confocal microscopy images of immunofluorescence experiments confirming HDAC6 and VIM are essential for aggresome formation** *(related to Figure 1).*

A) Full confocal microscope images of immunofluorescence experiments in unstressed or stressed VIM KO cells treated for 18 hours with MG132. VIM antibody staining confirmed the complete VIM protein knockout and Ub antibody staining showed impairment of aggresome formation in VIM KO cells. Scale bar is 20μm.

B) Full confocal microscope images of immunofluorescence experiments in unstressed or stressed VIM KO cells for 18 hours with MG132. No HDAC6+ and Ub+ aggregates were detected in VIM KO HeLa cells upon 18 hours proteasome inhibition. Scale bar is 20μm.

C) Full confocal microscope images of immunofluorescence experiments in unstressed or stressed HDAC6 KO cells for 18 hours with MG132. HDAC6 antibody staining confirmed the complete HDAC6 protein knockout and Ub antibody staining showed impairment of aggresome formation in HDAC6 KO cells. Scale bar is 20μm.

D) Full confocal microscope images of immunofluorescence experiments in unstressed or stressed HDAC6 KO cells for 18 hours with MG132. No VIM+ and Ub+ aggregates were detected in HDAC6 KO HeLa cells upon 18 hours proteasome inhibition. Scale bar is 20μm.

Fig.S3.

A. Aggresome biogenesis time points

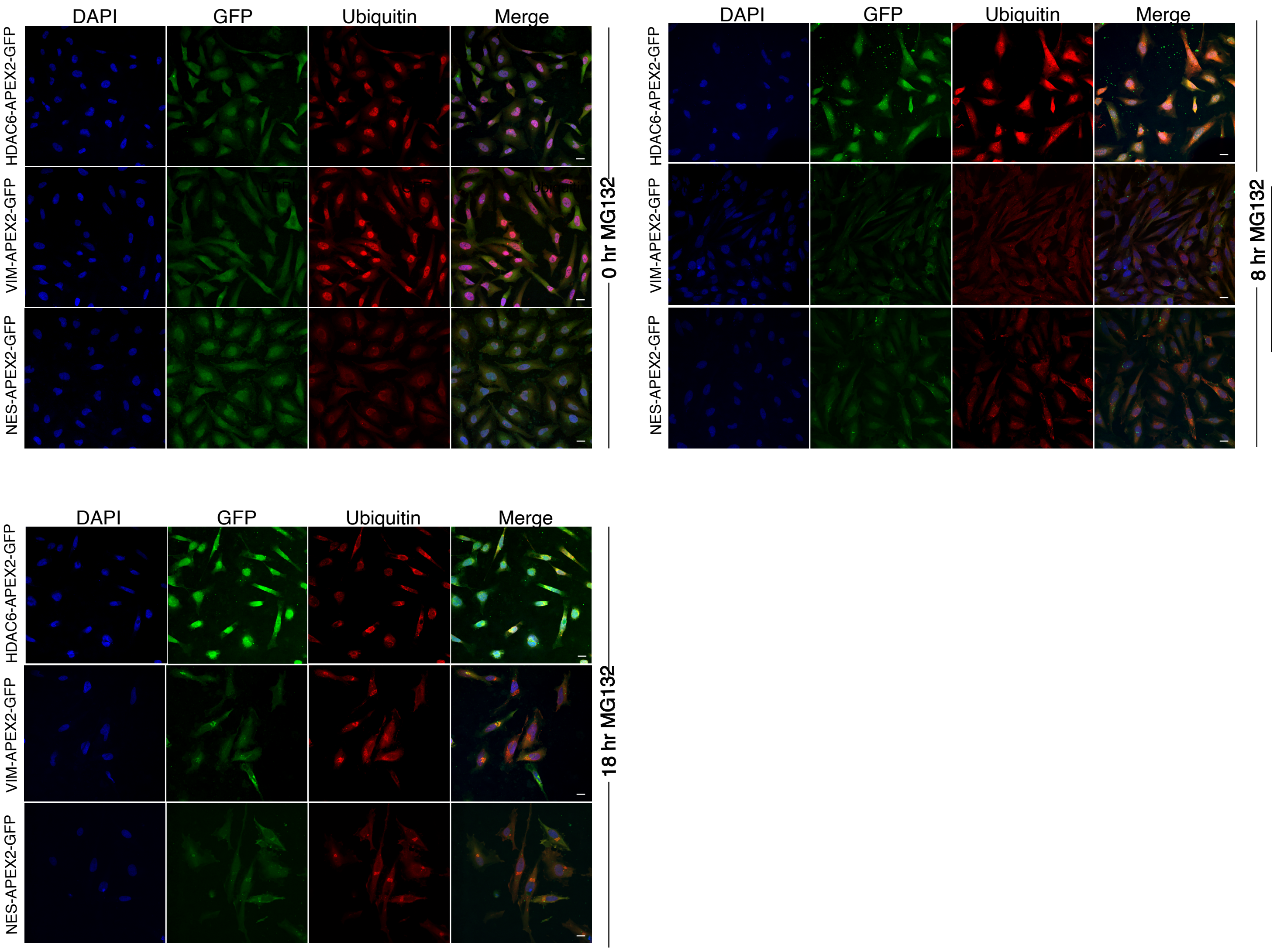


Aggresome clearance time points

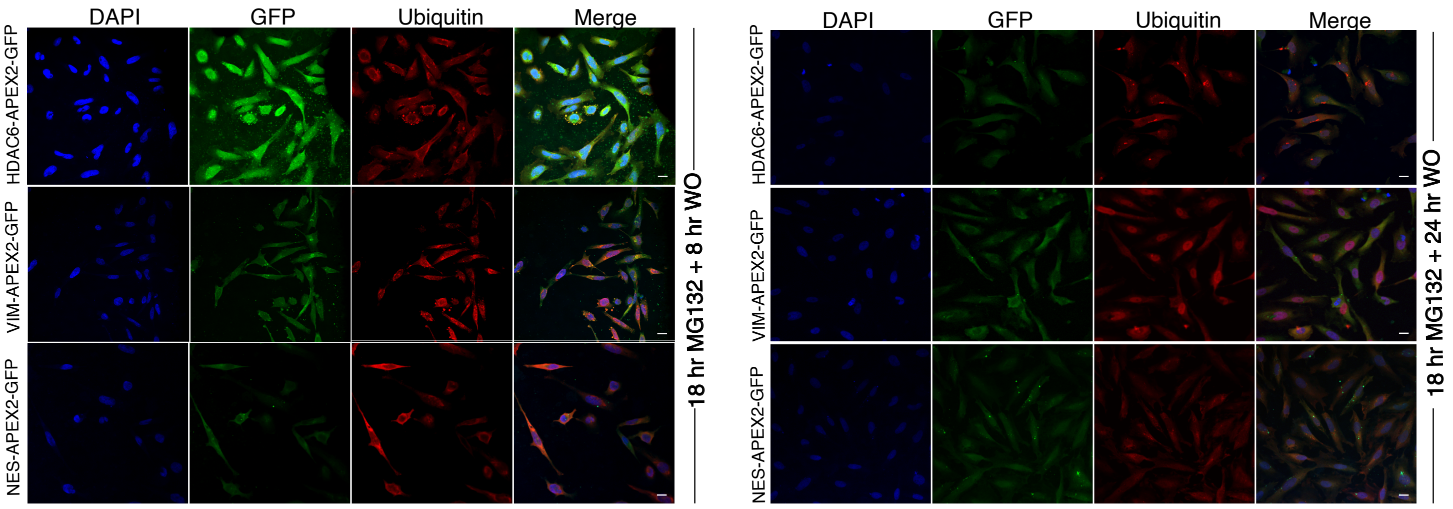

**Figure S3: Full confocal microscopy images of immunofluorescence experiments to detect different stages of APP** *(related to Figure 2).*

A) Full confocal microscope images of immunofluorescence experiments from HDAC6-APEX2-GFP, VIM-APEX2-GFP HeLa cells and NES-APEX2-GFP cells at different time points of MG132 stress exposure and washout. Cell lines were untreated or treated with MG132 for 8 hours, 18 hours, or treated for 18 hours followed by 8 hours or 24 hours of MG132 washout. Immunostaining was performed using HDAC6, VIM and Ub antibodies for each condition. Aggresomes or small intermediate mifolded aggregates are confirmed by overlap of HDAC6 or VIM signal with Ub signal (merged signal is yellow), as Ub is a marker for protein aggregation during proteasomal inhbition. Scale bar is 20μm.

### Fig.S4

A.

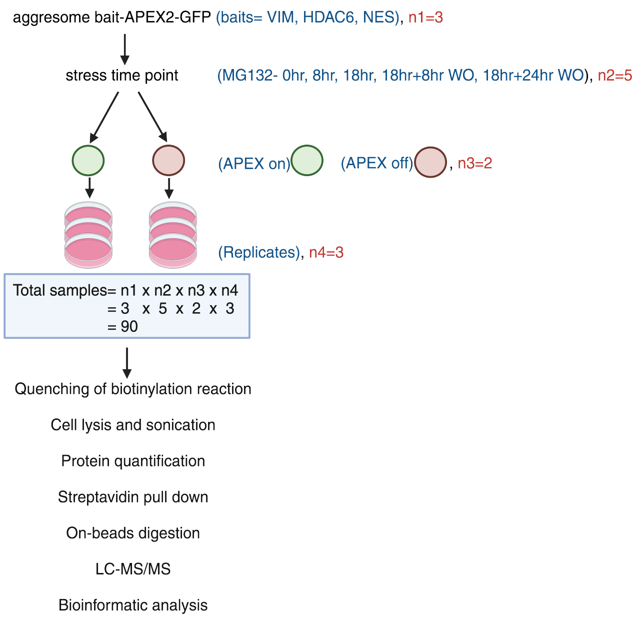


B.

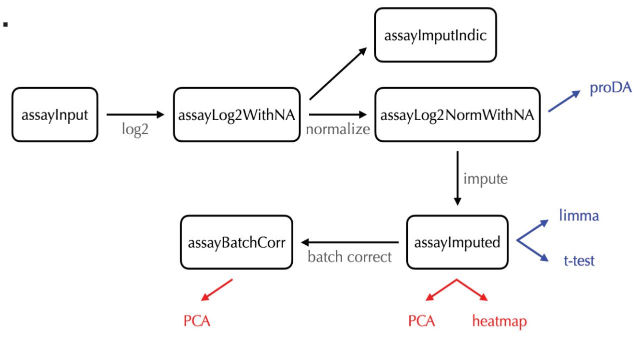


| | |
|---|---|
| assayInput | MaxLFQ.intensity |
| assayLog2WithNA | log2_MaxLFQ.intensity_withNA |
| assayImputIndic | imputed_MaxLFQ.intensity |
| assayLog2NormWithNA | log2_MaxLFQ.intensity_withNA_norm |
| assayImputed | log2_MaxLFQ.intensity_norm |
| assayBatchCorr | log2_MaxLFQ.intensity_norm |

C.

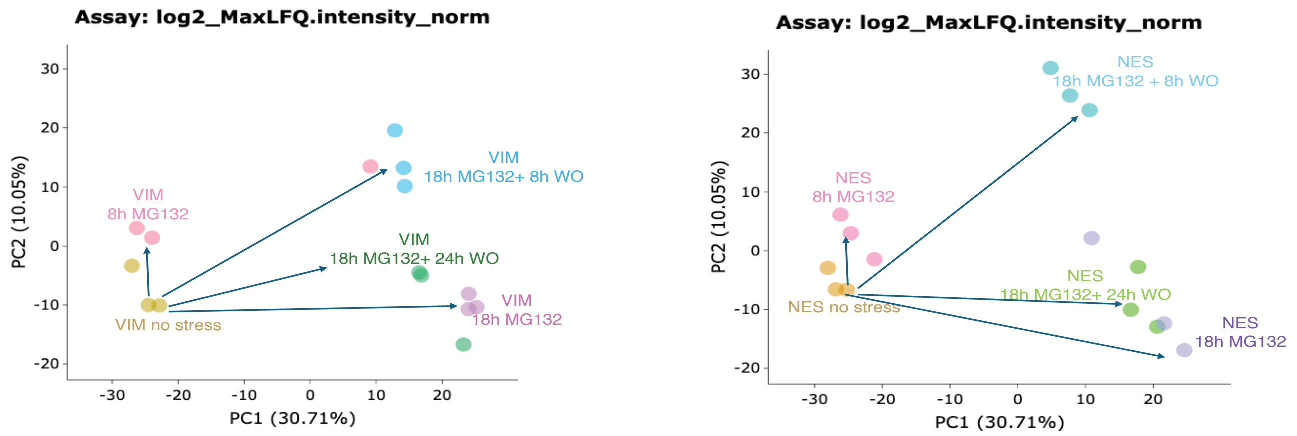


D.

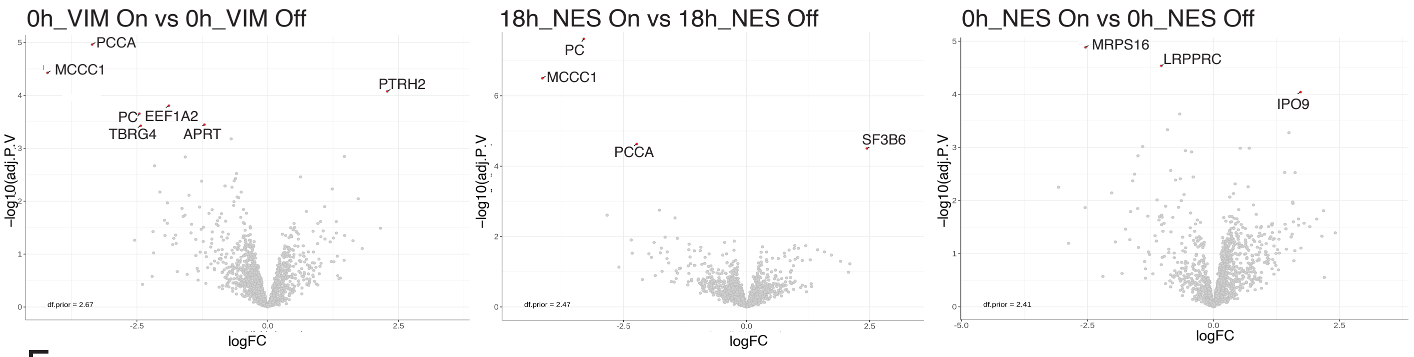


E.

Stress-specific proteins

| | | |
|---|---|---|
| A2M | EWSR1 | PPP6C |
| ABCB7 | FUS | PRAF2 |
| ABCD1 | G6PD | PRPS2 |
| ACP1 | GLRX | PRRC1 |
| ACSL4 | HCCS | PSME2 |
| AKAP2 | HMGB3 | RAB9A |
| AKR1A1 | HMGCS1 | RANBP1 |
| ANXA11 | HNRNPAB | RBM3 |
| ARMCX3 | ISG15 | RER1 |
| ATP6V1G1 | ITGA11 | RNH1 |
| ATXN2L | KCTD12 | RPS6KA3 |
| BAG3 | LAMP2 | SCARB2 |
| BANF1 | LASP1 | SCD |
| CALM1 | LPP | SEC13 |
| CLTA | MAP4 | SFN |
| CPPED1 | MAPRE2 | SMAD2 |
| CPT1A | NDUFAF2 | SMARCA2 |
| CSRP1 | NELFCD | SNW1 |
| CTTN | NKRF | SNX12 |
| DAZAP1 | NOSIP | TES |
| DDX19A | PABPC1 | TRA2B |
| DUSP1 | PDCD4 | TWF2 |
| EDIL3 | PDLIM1 | VAMP7 |
| EIF4A2 | PGM2L1 | XPNPEP1 |
| EIF4B | PHPT1 | |

F.

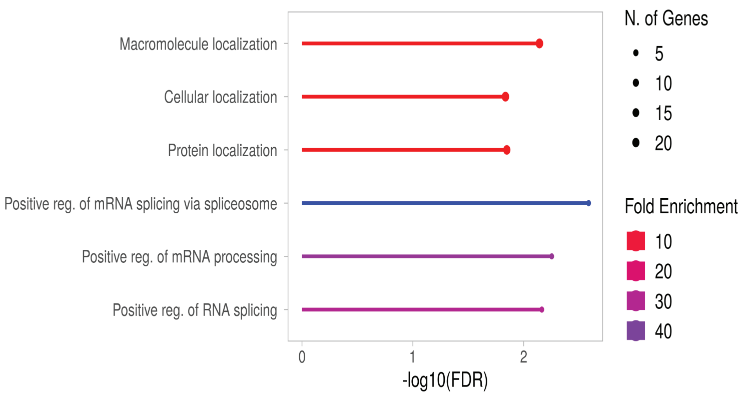

**Figure S4: Mass spectrometry analysis pipeline, data quality, and non-specific streptavidin binders in data of dual-bait APEX2 experiment (***related to Figure 3)*.

A) Analysis pipeline diagram of dual-bait APEX2 experiment (related to data shown in Figure 3, 4 and 5). Two aggresome marker proteins, HDAC6, VIM and one control, NES (all APEX2-GFP tagged) cell lines were treated with MG132 for various time points (0h, 8h, 18h, 18h+8h washout, 18h+24h washout), and APEX2 was either ON (BP, $H_2 0_2$) or OFF (only BP) for each condition. For each condition, we had 3 replicates. All samples were processed as described in the schematic and bioinformatics analysis was used to map the neighbourhood interaction networks of the bait proteins during the course of APP.

B) Overview of the workflow of LC-MS/MS analysis. In the table, the first column contains generic names representing the 'stage' that each assay corresponds to. The second column contains the actual assay names.

C) Principal component analysis (PCA) to obtain a reduced dimensionality representation of the data, in order to visualize the samples in two dimensions. Here, the first PCA analysis is a representation of the variance in LC-MS/MS data of all VIM-APEX2-GFP samples across all time points of stress. The second PCA analysis is a representation of the variance in LC-MS/MS data of all NES-APEX2-GFP samples across all time points of stress.

D) Volcano plots of the following comparisons as control: VIM APEX2 On versus APEX Off_0hr MG132, NES APEX2 On versus APEX Off_18hr MG132 and NES APEX2 On versus APEX Off_0hr MG132.

E) The list of stress-specific VIM-APEX2 hits at peak aggresome time point. Summation of upregulated proteins shared between comparison groups 1&3, 2&3 and 1,2&3.

F) Gene ontology analysis of biological function of the stress-specific VIM-APEX2 hits *(corresponding to Fig. 3E).*

Fig.S5

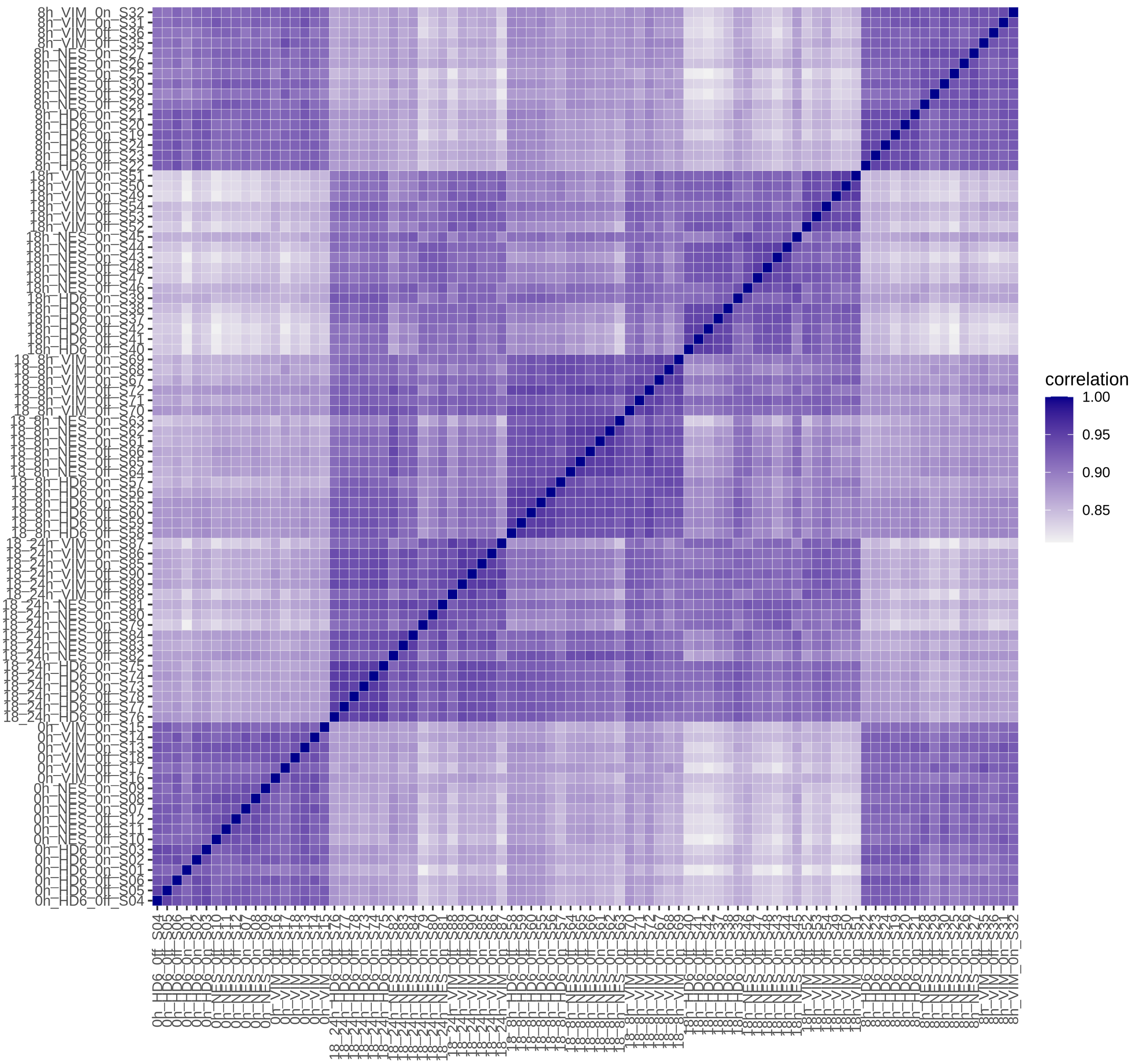


**Figure S5: Correlation plot of Pearson correlations between all pairs of samples (***related to Figure 3.***)**

The correlation plot was generated by Einprot. The plot shows the pairwise Pearson correlations between all pairs of samples, based on the log2_MaxLFQ.intensity_norm assay. The $\log_2$ MaxLFQ intensity norm assay refers to a normalized and $\log_2$-transformed version of MaxLFQ intensity values used in label-free quantification (LFQ) for mass spectrometry-based proteomics.

#### Fig.S6

A.

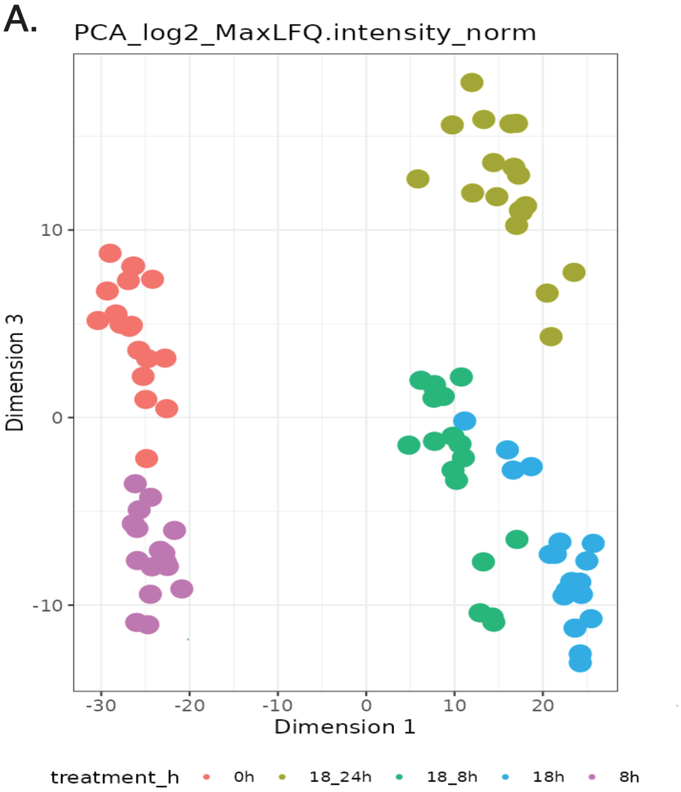


B.

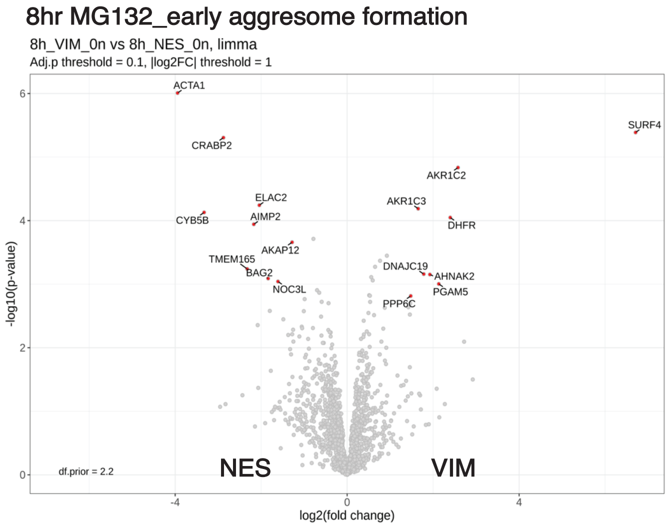


C.

##### 18+8 hr VIM apex on vs NES apex on
##### early aggresome clearance_GO Molecular Function

| N | High level GO category | Genes |
|---|---|---|
| 16 | GO:0036094 small molecule binding | PYGL ARF6 RAP2B SRPK1 RAB35 G6PD RAB8A AQR PRKACA GNA11 RAB23 CSNK1A1 RAB32 SAR1B MYO1E ACACA |
| 16 | GO:0097367 carbohydrate derivative binding | ARF6 RAP2B SRPK1 PYGL RAB35 RAB8A AQR PRKACA GNA11 RAB23 CSNK1A1 RAB32 HSD17B12 SAR1B MYO1E ACACA |
| 15 | GO:0016787 hydrolase activity | MTMR2 GNA11 RAB23 RAB32 SAR1B RNPEP RAP2B RAB35 MYO1E ARF6 RAB8A LAS1L AQR GBE1 TSN |
| 11 | GO:0016740 transferase activity | CSNK1A1 GBE1 SRPK1 PYGL SMS CPT1A PRKACA CPNE3 GTF3C4 CALM3 CALM1 |
| 11 | GO:0044877 protein-containing complex binding | GNA11 RBM3 MYO1E H1-4 PSMG1 NDUFAF2 NOC2L CUL2 HSD17B12 SIN3A RAB32 |
| 7 | GO:0098772 molecular function regulator activity | CALM3 CALM1 ARF6 SIN3A RPS27L RAP2B GTF3C4 |
| 6 | GO:0005215 transporter activity | SLC25A5 SFXN3 SLC38A2 TMEM165 SLC16A1 TMEM109 |
| 6 | GO:0022857 transmembrane transporter activity | SLC25A5 SFXN3 SLC38A2 TMEM165 SLC16A1 TMEM109 |
| 5 | GO:0016491 oxidoreductase activity | HSD17B12 CYP51A1 ALDH3A2 AKR1C2 G6PD |
| 4 | GO:0030234 enzyme regulator activity | CALM3 CALM1 RPS27L GTF3C4 |
| 3 | GO:0005198 structural molecule activity | RPS27L H1-4 KRT13 |
| 3 | GO:0008289 lipid binding | CPNE3 RAB35 MYO1E |

D.

##### 18+8 hr VIM apex on vs NES apex on
##### early aggresome clearance_GO Biological Process

Genes are grouped by functional categories defined by high-level GO terms.

| N | High level GO category | Genes |
|---|---|---|
| 25 | GO:0051234 establishment of localization | SLC25A5 SNAP23 AP1B1 SFXN3 RAB35 RAB23 RAB32 SLC38A2 TMEM165 SAR1B SLC16A1 MYO1E CALM3 ARF6 RAB8A TMED9 CALM1 PRKACA MTMR2 CPT1A TMEM109 COMMD9 G6PD NDUFAF2 CYP51A1 |
| 24 | GO:0065008 regulation of biological quality | SLC25A5 GNA11 SNAP23 TMEM165 AKR1C2 CALM3 RAB8A CNOT9 CALM1 PRKACA SAR1B RAP2B MTMR2 CPT1A COMMD9 HSD17B12 SLC16A1 NDUFAF2 ARF6 SIN3A ACACA PYGL NOP53 G6PD |
| 22 | GO:0044085 cellular component biogenesis | LAS1L SNAP23 SRPK1 NOP53 RAB23 HEATR1 PPAN SAR1B PSMG1 RPS27L NOC2L AP1B1 PRKACA RAB32 ARF6 RAB8A MTMR2 UTP20 NDUFAF2 H1-4 ACACA SIN3A |
| 20 | GO:0042221 response to chemical | CPNE3 GNA11 RAB8A SIN3A PRKACA AKR1C2 CALM3 CALM1 CRIP1 CPT1A RAB35 CNOT9 SLC16A1 G6PD NDUFAF2 ARF6 ALCAM ACACA NOP53 SLC38A2 |
| 19 | GO:0051641 cellular localization | SLC25A5 SNAP23 RAB35 RAB23 RAB32 SAR1B MYO1E CALM3 ARF6 RAB8A TMED9 CALM1 PRKACA MTMR2 AP1B1 NOP53 NDUFAF2 SIN3A CPT1A |
| 18 | GO:0032879 regulation of localization | SLC25A5 SAR1B CALM3 RAB8A RAP2B SIN3A CALM1 PRKACA MTMR2 CPT1A TMEM109 SLC16A1 G6PD NDUFAF2 ARF6 CYP51A1 CPNE3 RAB23 |
| 17 | GO:0023051 regulation of signaling | SLC25A5 CSNK1A1 S100A4 SIN3A AKR1C2 ARF6 PRKACA MTMR2 NOP53 CPT1A SLC16A1 CALM3 NDUFAF2 RAB8A CALM1 CNOT9 NOC2L |
| 17 | GO:0033036 macromolecule localization | RAB23 RAB32 SAR1B ARF6 RAB8A TMED9 AP1B1 PRKACA SNAP23 NOP53 CPT1A RAB35 SLC16A1 CALM3 NDUFAF2 SIN3A CYP51A1 |
| 16 | GO:0048583 regulation of response to stimulus | SLC25A5 CSNK1A1 S100A4 SIN3A AKR1C2 ARF6 PRKACA MTMR2 NOP53 CALM3 G6PD NDUFAF2 CALM1 SLC38A2 CNOT9 NOC2L |
| 14 | GO:0009893 positive regulation of metabolic process | RBM3 HEATR1 CNOT9 CSNK1A1 CEBPZ CALM3 RPS27L CALM1 NOP53 CPT1A RAP2B SIN3A SLC38A2 SLC25A5 |
| 13 | GO:0006950 response to stress | TMEM109 CRIP1 SRPK1 PRKACA RAP2B RPS27L SLC38A2 SIN3A SNAP23 NOP53 RAB23 G6PD KRT13 |
| 5 | **GO:0006914 autophagy** | **RAB23 CALM1 RAB8A SLC25A5 PRKACA** |

**Figure S6: VIM-APEX2 reveals time-dependent APP proteins, data quality and list of gene groups enriched in early aggresome clearance time point** (*related to Figure 4).*

A) PCA analysis representation of the variance in LC-MS/MS data for VIM, HDAC6 and NES-APEX2-GFP across each time points of stress.

B) Volcano plot showing statistically significant (adjusted p value < 0.05) VIM interactors in the following comparison: VIM APEX2 On versus NES APEX2 On at 8hr MG132 treatment (early aggresome formation).

C) Gene Ontology analysis of Molecular function of proteins enriched as VIM-APEX2 hits during early time point of aggresome clearance. The table groups the genes in molecular function categories ranked in order of the highest number of genes corresponding to a function. This was generated using ShinyGO 0.80.

D) Gene Ontology analysis of biological process of proteins enriched as VIM-APEX2 hits during early time point of aggresome clearance. The table groups the genes in biological process categories ranked in order of the highest number of genes corresponding to a process. This was generated using ShinyGO 0.80.

#### Fig.S7

### A.

18hr MG132 + 24hr wo_**HDAC6 vs NES_peak aggresome clearance time point**

GO Biological Process

Genes are grouped by functional categories defined by high-level GO terms.

| N | High level GO category | Genes |
|---|---|---|
| 11 | GO:0006950 response to stress | RPA2 TPR ACAA2 NPM1 ISG15 RFC1 SMC3 ACADM NIBAN1 PSIP1 TIAL1 |
| 11 | GO:0042221 response to chemical | TPR AKR1C2 ACAA2 NPM1 ISG15 ASAH1 KRT18 SRSF7 MKKS IMPDH2 MPST |
| 11 | GO:0051234 establishment of localization | TPR PMPCB MKKS NPM1 NUP210 KRT18 CCDC51 ACAA2 SRSF7 ATP5PO AFG3L2 |
| 10 | GO:0051641 cellular localization | TPR PMPCB MKKS NPM1 NUP210 KRT18 RPA2 ACAA2 ISG15 IK |
| 9 | GO:0009056 catabolic process | ACADM CLPX ACAA2 ISG15 MCCC1 ASAH1 UROD NPM1 MPST |

### B.

18hr MG132 + 24hr wo_**VIM vs NES_peak aggresome clearance time point**

GO Biological Process

Genes are grouped by functional categories defined by high-level GO terms.

| N | High level GO category | Genes |
|---|---|---|
| 10 | GO:0051234 establishment of localization | PMPCB MYO1E NUP210 ACSL4 ACAA2 SPTBN2 SFN APOO PITRM1 TUBA1C |
| 8 | GO:0009056 catabolic process | EXOG CLPX ACAA2 ISG15 SUCLG2 PCCA AKR1C3 MPST |
| 8 | GO:0044085 cellular component biogenesis | HCCS NDUFS8 HIGD2A PSIP1 ISG15 SPTBN2 NDUFB7 RCC1 |
| 8 | GO:0051641 cellular localization | PMPCB MYO1E NUP210 ACAA2 ISG15 SFN PITRM1 TUBA1C |
| 7 | GO:0033036 macromolecule localization | PMPCB SFN ACSL4 ISG15 NUP210 APOO PITRM1 |
| 7 | GO:0065008 regulation of biological quality | AKR1C2 AKR1C3 ACAA2 SPTBN2 SFN ISG15 ACSL4 |
| 6 | GO:0006950 response to stress | NDUFS8 ACAA2 SFN ISG15 AKR1C3 PSIP1 |

**Figure S7: List of gene groups enriched in dual-bait proximity labelling dataset at peak aggresome clearance time point-** *related to Figure 5.*

A) Gene Ontology analysis of biological process of proteins enriched as HDAC6-APEX2 hits during 18hr MG132+ 24hr washout time point, that is, during complete aggresome clearance. The table groups the genes in biological process categories ranked in order of the highest number of genes corresponding to a process. This was generated using ShinyGO 0.80.

B) Gene Ontology analysis of biological process of proteins enriched as VIM-APEX2 hits during 18hr MG132+ 24hr washout time point, that is, during complete aggresome clearance. The table groups the genes in biological process categories ranked in order of
